# Centromere identity is dependent on nuclear spatial organization

**DOI:** 10.1101/2021.12.16.473016

**Authors:** Weifang Wu, Toni McHugh, David A. Kelly, Alison L. Pidoux, Robin C. Allshire

## Abstract

The establishment of centromere-specific CENP-A chromatin is influenced by epigenetic and genetic processes. Central domain sequences from fission yeast centromeres are preferred substrates for CENP-A^Cnp1^ incorporation, but their use is context dependent, requiring adjacent heterochromatin. CENP-A^Cnp1^ overexpression bypasses heterochromatin dependency, suggesting heterochromatin ensures exposure to conditions or locations permissive for CENP-A^Cnp1^ assembly. Centromeres cluster around spindle-pole bodies (SPBs). We show that heterochromatin-bearing minichromosomes localize close to SPBs, consistent with this location promoting CENP-A^Cnp1^ incorporation. We demonstrate that heterochromatin-independent *de novo* CENP-A^Cnp1^ chromatin assembly occurs when central domain DNA is placed near, but not far from, endogenous centromeres or neocentromeres. Moreover, direct tethering of central domain DNA at SPBs permits CENP-A^Cnp1^ assembly, suggesting that the nuclear compartment surrounding SPBs is permissive for CENP-A^Cnp1^ incorporation because target sequences are exposed to high levels of CENP-A^Cnp1^ and associated assembly factors. Thus, nuclear spatial organization is a key epigenetic factor that influences centromere identity.

## Introduction

Centromeres are specialized chromosomal sites where multiprotein complexes known as kinetochores are assembled. Kinetochores attach chromosomes to spindle microtubules to mediate accurate mitotic and meiotic chromosome segregation. The assembly of kinetochores in many eukaryotes including yeasts and humans relies on specialized centromeric chromatin in which canonical histone H3 is replaced by the CENP-A/cenH3 histone H3 variant (Cnp1 in fission yeast, *Schizosaccharomyces pombe*). CENP-A-containing chromatin provides the underlying epigenetic mark that specifies the chromosomal site at which kinetochores assemble. CENP-A is required to establish and maintain centromere identity and thus indicates active centromeres (Allshire and Karpen, 2008; Mellone and Fachinetti, 2021).

In organisms with monocentric chromosomes centromeres are confined to a single locus on each chromosome. Such centromeres are often composed of long tandem arrays of repetitive sequences such as *α*-satellite repeats on human chromosomes (DeBose-Scarlett and Sullivan, 2021). These repeats provide a substrate for the *de novo* establishment of CENP-A chromatin and assembly of functional kinetochores when introduced into human cells. Thus, *α*-satellite repeats trigger centromere formation. Acentric chromosomes lacking centromeres are unable to attach to spindle microtubules and are lost during cell division. However, following centromere ablation through centromere inactivation or deletion of centromere DNA, neocentromeres can arise spontaneously at unusual locations that lack sequence similarity to normal centromere DNA but allow stable segregation of such acentric chromosomes (DeBose-Scarlett and Sullivan, 2021; Ishii et al., 2008). Thus, centromeric DNAs are not the only sequences that can trigger the assembly of functional kinetochores. Once assembled at a particular location, including neocentromeres or sites that do not usually incorporate CENP-A, CENP-A chromatin is stably propagated at that site though cell division using intrinsic maintenance mechanisms (Mitra et al., 2020; Westhorpe and Straight, 2015). Consequently, prior CENP-A assembly can mark a chromosomal locus for continued persistence of CENP-A chromatin on one homologue whereas the same locus remains devoid of CENP-A on the other (DeBose-Scarlett and Sullivan, 2021).

The fission yeast genome is carried on three monocentric chromosomes with regional centromeres of 40-110 kb comprised of two distinct domains (see Figure S1): CENP-A^Cnp1^ chromatin assembles across the central domain consisting of central core (*cc*) and flanking innermost repeat (*imr*) DNA, which are surrounded outer repeats (*otr-dg/dh*) assembled in Clr4 Histone H3 lysine 9 methyl-(H3K9me)-transferase-dependent heterochromatin (Allshire and Ekwall, 2015; Martienssen and Moazed, 2015). The central core of centromere 2 (*cc2*) is unique but the central cores of *cen1* and *cen3* share the same sequence. *imr* elements are unique to each centromere and mark the transition between CENP-A^Cnp1^ chromatin and the heterochromatic *otr-dg/dh* repeats which are conserved in sequence, but not arrangement, between the three centromeres (Clarke et al., 1993; Takahashi et al., 1992). tRNA genes that reside in each *imr* element demarcate these distinct centromeric domains and prevent heterochromatin from encroaching into the central CENP-A^Cnp1^ chromatin domain (Noma et al., 2006; Scott et al., 2006). Two divergent *Schizosaccharomyces* species (*S. octosporus* and *S. cryophilus*) share a similar centromere domain organization (Tong et al., 2019).

Like human *α*-satellite centromeric DNA, fission yeast central domain DNA is a preferred substrate for CENP-A^Cnp1^ and kinetochore assembly. This preferred status is underscored by the observation that, in contrast to other sequences, naïve central domain DNA borne on minichromosomes readily assembles and maintains CENP-A^Cnp1^ chromatin following transient CENP-A^Cnp1^ overexpression, bypassing the usual requirement for adjacent heterochromatin (Castillo et al., 2013; Catania et al., 2015). Interestingly, despite having no sequence homology with *S. pombe* centromeres, central domains from *S. octosporus* and *S. cryophilus* are competent to assemble CENP-A^Cnp1^ chromatin and functional centromeres in *S. pombe,* indicating that fission yeast central domains possess conserved instructive features (Tong et al., 2019). *S. pombe* central domain sequences are transcribed by RNAPII and exhibit high rates of histone H3 turnover which may contribute to the replacement of S phase-deposited placeholder H3 with CENP-A^Cnp1^ during the subsequent G2 (Shukla et al., 2018; Singh et al., 2020). H3 is evicted from central domain chromatin even in the absence of CENP-A and kinetochore proteins (Shukla et al., 2018). The Mis18 complex acts in concert with the CENP-A chaperone HJURP to recognize pre-existing CENP-A nucleosomes and ensure their persistence at particular locations by mediating H3 replacement with CENP-A in new H3-containing nucleosomes assembled during the preceding S-phase (Mitra et al., 2020; Westhorpe and Straight, 2015; Zasadzińska and Foltz, 2017). Thus, fission yeast central domain DNA possesses innate sequence-driven properties that program H3 eviction, making it a favored substrate for CENP-A^Cnp1^ chromatin assembly, which, once assembled is rendered heritable though an intricate read-write mechanism.

Centromeres are tightly clustered around spindle pole bodies (SPBs; centrosome equivalents) during interphase in both fission (*S. pombe)* and budding (*Saccharomyces cerevisiae)* yeast (Funabiki et al., 1993; Muller et al., 2019; Winey and O’Toole, 2001). In *S. cerevisiae* SPB-to-centromere microtubules persist in G1 and mediate SPB-centromere clustering (Guacci et al., 1997; Jaspersen, 2021; Winey and O’Toole, 2001). Proper centromere clustering around *S. pombe* SPBs is dependent on the functions of SPB component Sad1 (LINC complex SUN domain protein) and Lem2 (LEM domain inner nuclear membrane protein) which is distributed around the entire nuclear envelope (NE) but is concentrated at SPBs (Ebrahimi et al., 2018; Fernández-Álvarez et al., 2016). Csi1, resides at the kinetochore-SPB interface and is required for Lem2 accumulation around SPBs and acts with Lem2 to maintain SPB-centromere associations (Barrales et al., 2016; Ebrahimi et al., 2018; Hou et al., 2012). The CENP-A assembly factors Scm3^HJURP^, Mis16^RbAP46/48^, Mis18, Eic1/Mis19 are concentrated at centromeres clustered close to SPBs from late anaphase to prophase, including during G2 when new CENP-A^Cnp1^ is incorporated (Hayashi et al., 2004; Pidoux et al., 2009; Shukla et al., 2018; Subramanian et al., 2014; Williams et al., 2009).

Although fission yeast centromeric central domains are the preferred substrate for CENP-A^Cnp1^ assembly, establishment of CENP-A^Cnp1^ chromatin is subject to epigenetic regulation. The *de novo* assembly of CENP-A^Cnp1^ chromatin and functional centromeres on central domain sequences is dependent on the presence of adjacent outer repeat heterochromatin (see Figure S2) (Folco et al., 2008; Steiner and Clarke, 1994). Direct transformation of naked minichromosomes into cells lacking heterochromatin compared to crossing preassembled minichromosomes from wild-type cells results in a different fate: in the former central domain is assembled in H3 chromatin, in the latter it is assembled in CENP-A ^Cnp1^ chromatin (see Figure S2B, C) (Folco et al., 2008). These observations indicate that both context and prior history are important for determining chromatin state. Synthetic heterochromatin, assembled by tethering the Clr4 H3K9-methyltransferase, substitutes for outer repeats in promoting CENP-A^Cnp1^ assembly on minichromosomes when placed next to central domain DNA (see Figure S2F) (Kagansky et al., 2009). Thus, the properties of adjacent heterochromatin itself rather than other features of outer repeat elements, are critical for *de novo* CENP-A^Cnp1^ assembly. Heterochromatin could promote the establishment of CENP-A^Cnp1^ chromatin by recruitment of chromatin modifiers that influence turnover or other properties of histone H3 chromatin on adjacent central domain to favor CENP-A^Cnp1^ deposition (Catania et al., 2015; Shukla et al., 2018). Alternatively, since CENP-A^Cnp1^ overexpression circumvents the need for flanking heterochromatin in such CENP-A^Cnp1^ chromatin establishment assays (Catania et al., 2015) (see Figure S2) and endogenous heterochromatin domains are located at the nuclear periphery (Alfredsson-Timmins et al., 2007; Funabiki et al., 1993; Pichugina et al., 2016), it is possible that centromeric heterochromatin places such minichromosomes at a nuclear location that encourages *de novo* CENP-A^Cnp1^ chromatin assembly. The centromere clusters at *S. pombe* SPBs would be expected to provide a compartment naturally enriched with CENP-A^Cnp1^ and its loading factors.

Here we test if the positioning of centromeric DNA relative to existing centromeres and/or SPBs influences *de novo* CENP-A^Cnp1^ chromatin assembly and recruitment of kinetochore proteins. Heterochromatin-bearing plasmids localize close to SPBs, suggesting that heterochromatin may play a positioning role in promoting establishment of CENP-A^Cnp1^ chromatin. We demonstrate that potentially functional centromeric central domain DNA does not assemble CENP-A^Cnp1^ or kinetochore proteins unless inserted close to an already functional native centromere or neocentromere. Thus, proximity to an existing centromere *in cis* on the same chromosome promotes CENP-A^Cnp1^ and kinetochore assembly. Direct tethering of naïve minichromosome-borne central domain DNA to SPB-associated proteins in the absence of flanking heterochromatin revealed that proximity *in trans* to SPB-centromere clusters is also sufficient to trigger assembly of CENP-A^Cnp1^ chromatin and recruitment of kinetochore components. Thus, we define a key role for spatial genome organization, in particular centromere clustering, and resulting nuclear compartmentalization in determining centromere identity. Our findings reveal that centromeric heterochromatin functions to position centromeres within a nuclear compartment that ensures *de novo* CENP-A^Cnp1^ chromatin assembly.

## Results

### Centromeric heterochromatin mediates association with the SPB-centromere cluster

The *de novo* assembly of CENP-A^Cnp1^ chromatin on naïve centromeric central domain DNA freshly introduced into fission yeast as DNA by transformation on plasmid-based minichromosomes requires H3K9me-dependent heterochromatin formation on flanking outer dg/dh (K/L) centromere repeat DNA (Figure S2A, B and D) (Folco et al., 2008). Heterochromatin may influence CENP-A^Cnp1^ chromatin establishment through nuclear positioning cues. To test if centromeric heterochromatin promotes association with SPBs we utilized autonomously replicating minichromosomes which are less constrained than endogenous chromosomal regions with respect to their positioning within nuclei. In all strains used, 6 kb of endogenous *cc2* was replaced with 5.5 kb of *cen1* central domain DNA (*cc2*Δ*::cc1*; Figure S1B). Thus, *cc2* DNA carried by minichromosomes are unique sequences in these strains, allowing their specific analysis by quantitative chromatin immunoprecipitation (qChIP). As a consequence of this manipulation, sequences common to wild-type *cc1* and *cc3* are present at all three endogenous centromeres in *cc2*Δ*::cc1* cells and provide a positive control comparator for CENP-A^Cnp1^ and kinetochore protein association. The pHet minichromosome carries outer repeat DNA (*K”,* 2 kb) that is sufficient to trigger Clr4-dependent *de novo* heterochromatin formation when transformed into wild-type, but not *clr4*Δ, cells (Figure 1A, B; Table S2) (Folco et al., 2008; Kagansky et al., 2009). pcc2 carries 8.6 kb of *cen2* central domain DNA but lacks outer repeat DNA (Figure 1A; Table S2) and thus cannot form heterochromatin and does not assemble CENP-A^Cnp1^ chromatin or kinetochores (Figure 1C; Figure S2D) (Catania et al., 2015; Folco et al., 2008). However, pHcc2, carrying both outer repeat and *cc2* DNA (Figure 1A; Table S2) forms heterochromatin which permits CENP-A^Cnp1^ chromatin (Figure 1C, D), kinetochores and functional centromeres to be frequently established in wild-type cells following transformation (Figure S2A, Figure S3) (Catania et al., 2015; Folco et al., 2008).

**Figure 1.**
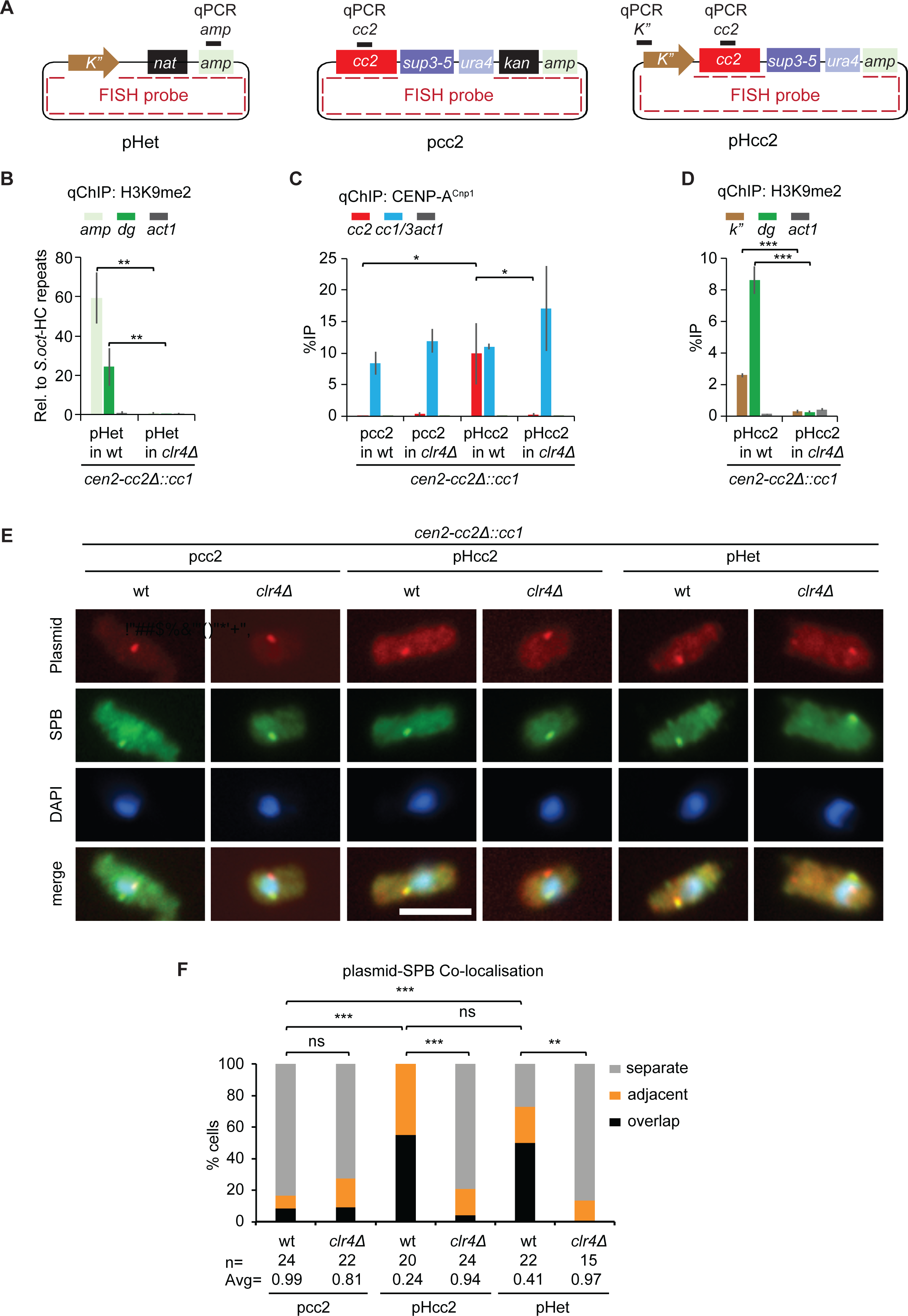
Centromeric heterochromatin associates with the SPB-centromere cluster. (A) Diagram of pHet, pcc2 and pHcc2 minichromosomes. Black bars above each plasmid map represent qChIP primer sites on ampicillin gene (*amp*), *cc2* and *K”* repeats of plasmids, respectively. Dashed red line on plasmids indicates position of FISH probe. (B-D) qChIP analyses for H3K9me2 levels on *amp* gene of pHet (B), *K”* repeats of pHcc2 (D), *dg* repeats of centromeric heterochromatin and *act1* gene (B, D), CENP-A^Cnp1^ levels on *cc2*, *cc1/3* (indicates sequences common to *cc1* and *cc3*) and *act1* in wt and *clr4*Δ cells containing *cc2*Δ::*cc1* at *cen2* transformed with pHet (B), pcc2 (C) or pHcc2 (C, D). %IP levels in *S. pombe* were normalized to %IP of *cen3* heterochromatin (HC) repeats from spiked-in *S. octosporus* chromatin in (B). qChIP results in (C, D) are reported as %IP. Data are mean ± SD (error bars) (n=3-4 experimental replicates). *, p<0.05; **, p<0.005; ***, p<0.0005 (Unpaired t-test). (E) Representative images of plasmid DNA FISH (red; probe as indicated in A), SPB location (green; anti-Cdc11) and DNA staining (blue, DAPI) in wt and *clr4*Δ cells transformed with pcc2, pHcc2 or pHet. Images were scaled relative to the maximum values of histogram. Scale bar, 5 μm. (F) Cells were classified into three groups according to the 3D distances between plasmid and SPB (Cdc11): overlap (≤0.3 μm), adjacent (0.3-0.5 μm) or separate (0.5-3 μm). Percentage of interphase cells (n, number analyzed from 3 independent experiments) in each category. Avg, average distance. ns, no significance; **, p<0.001; ***, p < 0.0001 (Mann-Whitney U test) (See also STAR Methods and Figures S1-S3).

Fluorescent In Situ Hybridization (FISH) to the backbone plasmid and/or cc2 sequences allowed pHet, pcc2 or pHcc2 minichromosome localization in wild-type cells relative to SPBs (Cdc11, SPB- specific centriolin ortholog; Figure 1A). pcc2 was found at, or in close proximity to, SPBs in 17% of cells, however, the presence of a heterochromatic repeat on pHcc2 with resulting CENP-A^Cnp1^ and kinetochore assembly increased SPB association to 100% (Figure 1E, F). Consistent with a requirement for heterochromatin for SPB association, only low levels of pHcc2-SPB association were detected in *clr4*Δ cells where heterochromatin and CENP-A^Cnp1^/kinetochores are unable to assemble (Figures 1C-F; S3A). Moreover, pHet, which only assembles heterochromatin, localized close to SPBs in 73% of wild type cells but only 13% of *clr4*Δ cells (Figure 1B, E and F).

Together these data indicate that centromeric outer repeat-induced heterochromatin is sufficient to mediate frequent contact with SPBs where centromeres and CENP-A^Cnp1^ assembly factors are concentrated. Thus, we propose that centromeric heterochromatin promotes exposure of adjacent *cc2* centromere DNA to this CENP-A^Cnp1^ assembly factor-rich nuclear compartment, thereby ensuring the assembly of CENP-A^Cnp1^ chromatin and kinetochores.

### Centromeric central domain DNA assembles CENP-A^Cnp1^ chromatin when inserted close to native centromeres

To test if a nuclear compartment formed by SPB-centromere clustering might stimulate *de novo* CENP-A^Cnp1^ chromatin assembly we inserted 8.6 kb of *cc2* DNA near or far from *cen1* and assayed for the presence of CENP-A^Cnp1^ chromatin. In all strains used endogenous *cc2* had been replaced with *cc1* so that regions L-to-Q of the resulting 8.6 kb *cc2* insertions are unique (*cc2*Δ::*cc1*; Figure S1B). The *lys1* locus resides just 26 kb from *cc1*, 11.3 kb from the left *otr1* heterochromatin repeat while *ade3* is a distant 2,438 kb from *cc1* (Figure 2A). Microscopy measurements demonstrated that *lys1* and *ade3* decorated with LacI-GFP on *lacO*-array insertions (Ding and Hiraoka, 2017) are positioned in close proximity to or distant from SPBs, respectively, in three dimensional nuclear space (Figure 2B, C). qChIP analysis showed that CENP-A^Cnp1^ was uniformly incorporated onto regions L-P across *cc2* following insertion at *lys1* (*lys1:cc2*). In contrast, no CENP-A^Cnp1^ enrichment was observed on *cc2* inserted at *ade3* (*ade3:cc2*) (Figure 2D). In addition, kinetochore proteins CENP-C^Cnp3^, CENP-K^Sim4^ and Knl1^Spc7^ were also recruited to *lys1:cc2* (Figure 2E-G), indicating that CENP-A^Cnp1^ deposition on *lys1:cc2* results in recruitment of both inner and outer kinetochore proteins. CENP-A^Cnp1^ was also incorporated on *cc2* inserted at *sdh1*, 24 kb to the right of *cen1-cc1*, or at a location we named *itg10*, 27 kb from on the right side of *cen2-cc2*Δ::*cc1* (*itg10:cc2*; Figures S4A, B). Insertion of *cc2* at locations 41 kb (*vps29:cc2*) and 47 kb (*bud6:cc2*) further away on the left side of *cen1-cc1* resulted in progressively less CENP-A^Cnp1^ incorporation, suggesting that the level incorporated on inserted *cc2* DNA is dependent on its proximity *in cis* to functional *cen1* (Figure S4C).

**Figure 2.**
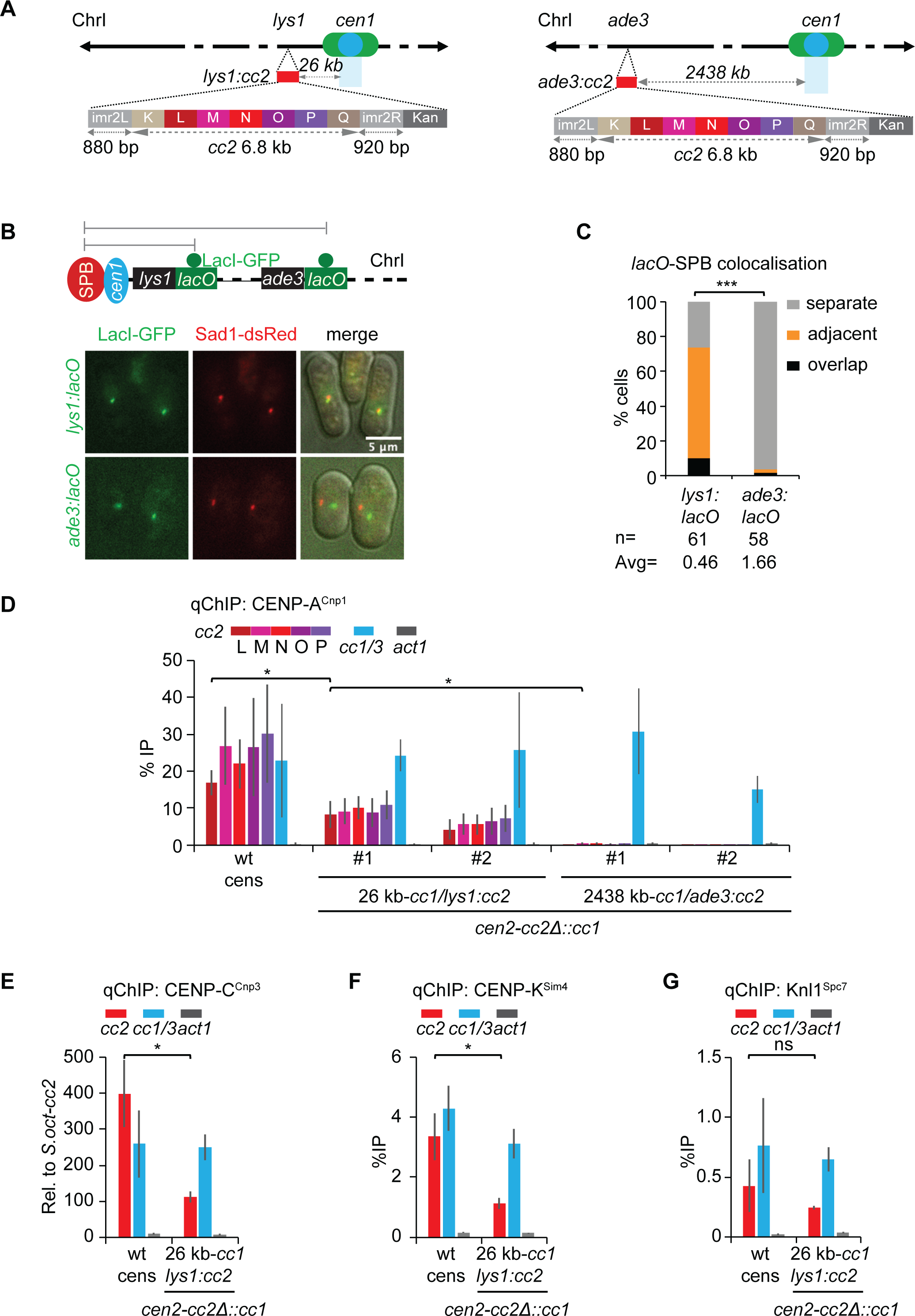
CENP-A^Cnp1^ chromatin is established on centromere-adjacent *lys1:cc2* central domain. (A) Ectopic *cc2*, carrying 880 bp *imr2L*, 6.8 kb *cc2* (subdivided into K-to-Q regions; 6 kb is unique) and 920 bp *imr2R* DNA, were inserted at *lys1* (*lys1:cc2*; 26 kb from *cc1*) or *ade3* (*ade3:cc2*; 2438 kb from *cc1*) on ChrI in *cc2*Δ::*cc1* strain. (B) Representative images of live cells expressing Sad1-dsRed (SPB marker) and LacI-GFP bound to *lys1:lacO* or *ade3:lacO* (Ding and Hiraoka, 2017). Images were scaled relative to the maximum intensity in the set of images. Scale bar, 5 μm. (C) 3D distances between *lys1:lacO* or *ade3:lacO* and SPBs (Sad1). Percentage of G2 cells (n, number analyzed from 3 independent experiments) in each category, classified as in Figure 1. Avg, average distance. ***, p < 0.0001 (Mann-Whitney U test) (See also STAR Methods.) (D) qChIP for CENP-A^Cnp1^ at regions L-P of *cc2, cc1/3* and *act1* in wt cens strain carrying endogenous *cen2-cc2* or *cen2-cc2*Δ::*cc1* strain with *lys1:cc2* or *ade3:cc2* insertions. # number indicates individual isolates. (E-G) qChIP analyses for CENP-C^Cnp3^ (E), CENP-K^Sim4^ (F), Knl1^Spc7^ (G) levels at *cc2, cc1/3* and *act1* gene in in wt cens strain carrying endogenous *cen2-cc2* or *cen2-cc2*Δ::*cc1* strain with *lys1:cc2*. %IP levels in *S. pombe* were normalized to %IP of *S. octosporus* central core from spiked-in chromatin in (E). qChIP results in (D, F, G) were reported as %IP. Data are mean ± SD (n=3). ns, no significance; *, p<0.05 (Unpaired t-test) (See also Figure S1, S4, S5).

Thus, either proximity to an endogenous centromere *in cis* on the same chromosome, or exposure to a distinct nuclear compartment formed by SPB-centromere clusters, effectively mediates *de novo* CENP-A^Cnp1^ assembly and kinetochore protein recruitment on naïve central domain DNA. *cc2* DNA inserted close to *cen1* might acquire CENP-A^Cnp1^ chromatin as a result of it spreading from *cen1* into *lys1:cc2*. However, little or no CENP-A^Cnp1^ enrichment was detected at three positions (i-iii) between *cen1* and *lys1:cc2* (Figure S4D). Thus CENP-A^Cnp1^ does not uniformly spread along the chromosome from its normal location at *cen1-cc1* into the *lys1:cc2* insert.

These analyses demonstrate that *cc2* DNA, a known substrate for fission yeast CENP-A^Cnp1^ and kinetochore assembly, incorporates CENP-A^Cnp1^ when inserted *in cis* close to native centromeres. The finding that the levels of CENP-A^Cnp1^ incorporated decrease with increasing distance from a centromere suggests that proximity to native centromeres provides an environment that is more favorable for CENP-A^Cnp1^ and kinetochore assembly on naïve centromere DNA.

### Proximity to functional centromeres, not locus-specific context, promotes CENP-A^Cnp1^ chromatin establishment

Neocentromeres form near fission yeast telomeres when an endogenous centromere is deleted (Ishii et al., 2008) (Figure 3A). Deletion of *cen1* (*cen1*Δ) results in neocentromeres being formed over the left (*neo1L; cd39*) or right (*neo1R; cd60*) subtelomeric regions on chromosome I (Ishii et al., 2008). FISH demonstrates that prior to neocentromere formation the subtelomeric *neo1R* locus is not located near SPBs whereas upon CENP-A^Cnp1^ assembly and neocentromere formation *neo1R* joins the interphase SPB-centromere cluster in 94% of cells, where CENP-A is concentrated (Figure 3A-E). Unlike when *cc2* was inserted at *lys1* in cells with a nearby functional *cen1* (Figure 2), insertion of *cc2* at *lys1* in *cen1*Δ cells with the *neo1R* neocentromere 1.8 Mb away failed to incorporate CENP-A^Cnp1^ (Figure 4A, B). This finding suggests that CENP-A^Cnp1^ fails to be incorporated at *lys1:cc2* upon insertion in cells with this neocentromere because *lys1* is displaced from the centromere cluster. Thus, a prediction is that insertion of *cc2* close to a region where neocentromeres can form will only result in CENP-A^Cnp1^ incorporation when an active neocentromere is present. We therefore inserted *cc2* at locations 73 kb (*itg6*), 60 kb (*itg7*) and 7 kb (*itg8*) from the *neo1R* region in cells with a wild-type *cen1* (no sub-telomeric neocentromere) or with an active neocentromere *neo1R* (wild-type *cen1* deleted) (Figure 4A).

**Figure 3.**
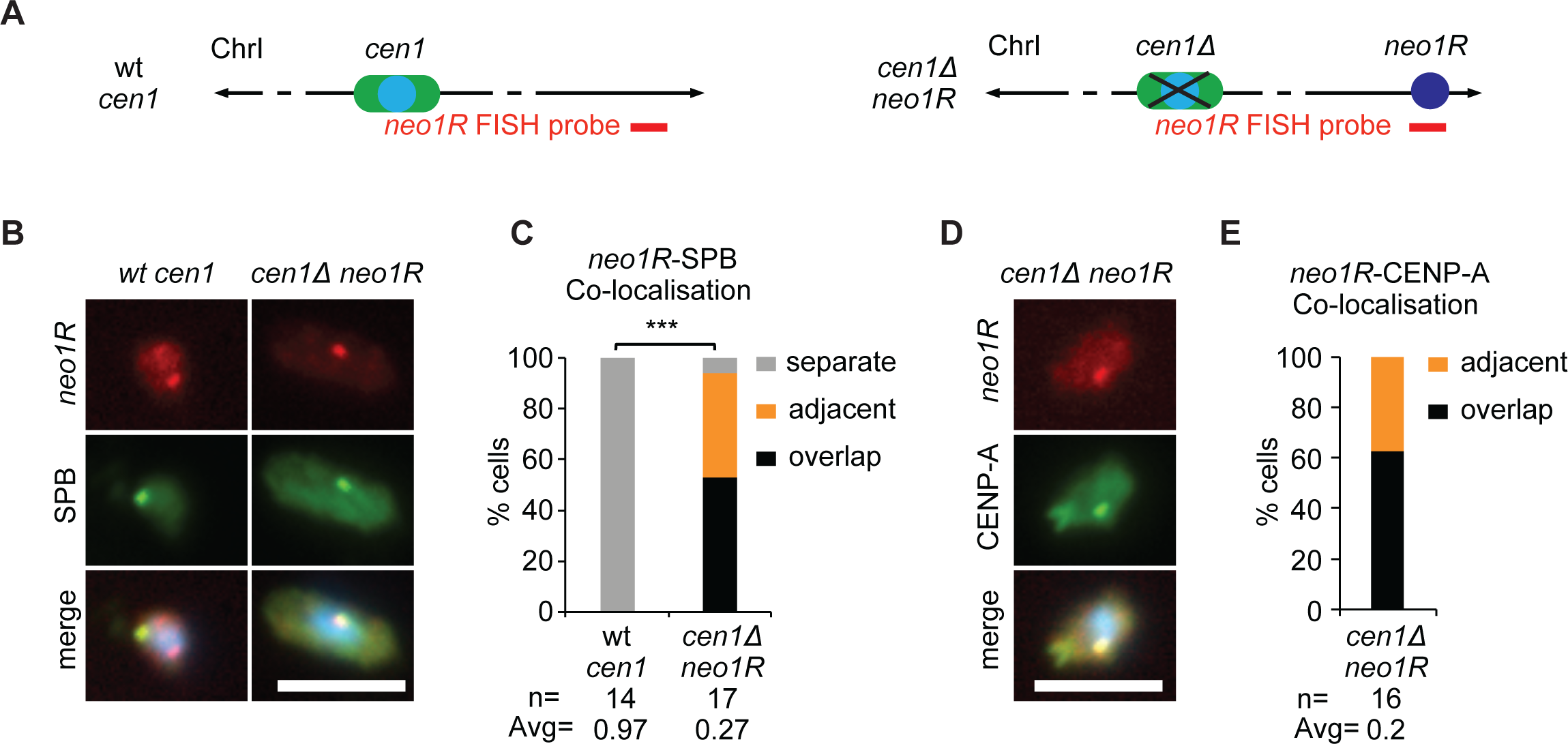
*neo1R* neocentromere clusters with endogenous centromeres at the SPB during interphase. (A) Diagram represents strains with *cen1* or lacking *cen1* but carrying *neo1R* neocentromere (*cen1*Δ *neo1R*). Red line indicates position of *neo1R* DNA FISH probe (ChrI: 5,513,871-5,530,124). (B, D) Representative images of *neo1R* DNA FISH (red; probe as indicated in A), SPB location (green; anti-Cdc11; B) or centromere clusters (green; anti-CENP^Cnp1^; D) and DNA staining (blue, DAPI) in wt cen1 (B) and *cen1*Δ *neo1R* cells. Images were scaled as in Figure 1. Scale bar, 5 μm. (C, E) 3D distances between *neo1R* DNA and SPBs (Cdc11; C) or centromere clusters (CENP-A^Cnp1^; E). Percentage of interphase cells (n, number analyzed) in each distance category, classified as in Figure 1. Avg, average distance. ***, p < 0.0001 (Mann-Whitney U test).

**Figure 4.**
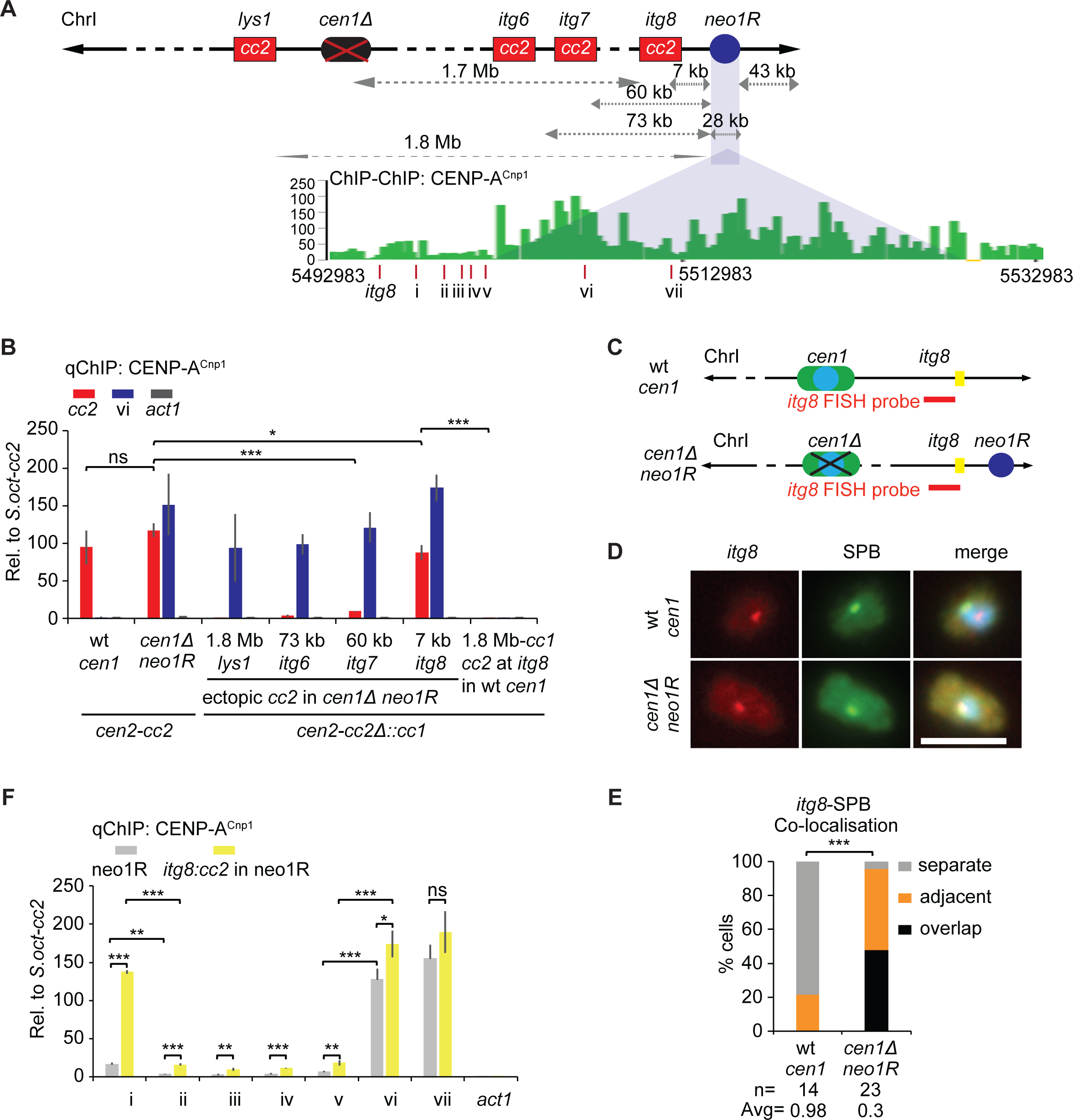
CENP-A^Cnp1^ chromatin can establish on the inserted neocentromere proximal-central domain. (A) Ectopic *cc2* inserted at *lys1*, *itg6* (ChrI: 5,435,010-5,435,237), *itg7* (ChrI: 5,447,816-5,448,235) and *itg8* (ChrI: 5,501,647-5,502,134), 1.8 Mb, 73 kb, 60 kb and 7 kb from *neo1R* CENP-A^Cnp1^ domain respectively. ChIP-CHIP analysis for CENP-A^Cnp1^ in *cen1*Δ *neo1R* (*cd60*) strain was obtained from (Ishii et al., 2008). Red lines indicate *itg8* and 7 qChIP primer sites (i-vii). (B) qChIP analyses of CENP-A^Cnp1^ levels at *cc2, cc1/3* and *act1* in wt *cen1* or *cen1*Δ *neo1R* strain with *lys1:cc2*, *itg6:cc2*, *itg7:cc2* or *itg8:cc2* insertions (genome positions as indicated in A). (C) Diagram represents wt-*cen1* or *cen1*Δ *neo1R* strains. Red line indicates position of *itg8* DNA FISH probe (ChrI 5,495,975-5,508,459). (D) Representative images of *itg8* DNA FISH (red; probe as indicated in C), SPB location (green; anti-Cdc11; B) and DNA staining (blue, DAPI) in wt *cen1* (B) and *cen1*Δ *neo1R* cells. Images scaled as in Figure 1. Scale bar, 5 μm. (E) 3D distances between *itg8* and SPBs (Cdc11), percentage of interphase cells (n, number analyzed) in each category, classified as in Figure 1. Avg, average distance. ***, p < 0.0001 (Mann-Whitney U test). (F) qChIP analyses for CENP-A^Cnp1^ levels at 7 loci (i-vii, positions as indicated in A) and *act1* in *cen1*Δ *neo1R* strain with or without *itg8:cc2* insertion. %IP levels in *S. pombe* were normalized to %IP of *S. octosporus* central core from spiked-in chromatin (B, F). Data are mean ± SD (n=3). ns, no significance; *, p<0.05; **, p<0.005; ***, p<0.0005 (Unpaired t-test).

Unlike wild-type cells where *itg8* was spatially distant from the SPB, *itg8* was positioned close to the SPB-centromere cluster in 96% of *neo1R* cells (Figure 4C-E). CENP-A^Cnp1^ was enriched on *lys1:cc2,* but not *itg8:cc2,* in cells with wild-type *cen1* (Figure 2D; Figure 4B). However, in cells with *cen1*Δ *neo1R* the pattern was reversed: no CENP-A^Cnp1^ incorporation occurred on *lys1:cc2* whereas high levels of CENP-A^Cnp1^ were detected on *itg8:cc2* located 8 kb from the active *neo1R* neocentromere. Little or no CENP-A^Cnp1^ was detected on the *itg6:cc2* and *itg7:cc2* insertions at greater distances from this neocentromere (Figure 4B). In cells with CENP-A^Cnp1^ incorporation at *itg8:cc2* we tested for CENP-A^Cnp1^ enrichment at five positions (sites i-v) between *itg8:cc2* and the active *neo1R* centromere (Figure 4A). As expected, high levels of CENP-A^Cnp1^ were detected at two positions (sites vi, vii) within the characterized *neo1R* neocentromere (Ishii et al., 2008). However, substantial CENP-A^Cnp1^ incorporation was only observed 5.2 kb from *neo1R* (site i; 1.6 kb from *itg8*) when *cc2* had been inserted at *itg8*, whereas little or no CENP-A^Cnp1^ enrichment was detected at sites i-v between *itg8* and *neo1R* (Figure 4F).

These analyses demonstrate that deletion of native *cen1* prevents *de novo* CENP-A^Cnp1^ incorporation on *cc2* subsequently inserted at *lys1* but permits CENP-A^Cnp1^ assembly on *cc2* when inserted close to a resulting neocentromere. The fact that CENP-A^Cnp1^ is not detected at most positions between the *neo1R* centromere and *itg8:cc2* indicates that, as at native *cen1 (*Figure S4D), CENP-A^Cnp1^ chromatin does not spread uniformly from the pre-existing neocentromere to the nearby inserted *cc2* DNA. We conclude that it is the proximity of *lys1* or *itg8* to functional centromeres, rather than properties of sequences immediately flanking these loci that allows the naturally CENP-A^Cnp1^-permissive *cc2* DNA substrate to assemble CENP-A^Cnp1^ when inserted at these locations.

### Centromeric heterochromatin is not required for *de novo* CENP-A^Cnp1^ incorporation on centromere DNA placed close to an existing centromere

In minichromosome-based establishment assays H3K9me-dependent heterochromatin is needed to allow *de novo* CENP-A^Cnp1^ incorporation on adjacent *cc2* central domain DNA (Figure S2) (Folco et al., 2008). If the nuclear environment formed by SPB-centromere clustering is sufficient to promote *de novo* CENP-A^Cnp1^ assembly, a prediction is that centromeric heterochromatin would not be required when central domain DNA is inserted close to endogenous centromeres. The *lys1*:*cc2* insertion is positioned only 11.3 kb from endogenous *cen1* heterochromatic *dh/otr1* repeats (Figure S5A). To determine if centromeric heterochromatin influences CENP-A^Cnp1^ chromatin establishment at *lys1* we inserted *cc2* DNA at this locus in either wild-type (wt) or heterochromatin-deficient *clr4*Δ cells (lack Clr4 H3K9 methyltransferase). FISH confirmed that the *lys1* locus and *lys1*:*cc2* insertion remain near SPBs in cells lacking Clr4 (Figure S5B-E). qChIP demonstrated that CENP-A^Cnp1^ was established on *lys1:cc2* insertions made in either wild-type or *clr4*Δ cells and that both CENP-C^Cnp3^ and Knl1^Spc7^ kinetochore proteins were recruited (Figure S5F-H). Thus, the *de novo* assembly of CENP-A^Cnp1^ and kinetochore proteins at *lys1:cc2* occurs independently of nearby centromeric heterochromatin.

We conclude that centromeric heterochromatin is not required to assemble CENP-A^Cnp1^ and kinetochore proteins on freshly introduced centromeric DNA if that DNA is positioned *in cis* close to an existing centromere which clusters with other centromeres, and associated CENP-A^Cnp1^ plus its assembly factors, around SPBs. The placement of centromeric central domain DNA close to active centromeres bypasses the requirement for heterochromatin. This lack of a need for centromeric heterochromatin is consistent with heterochromatin normally influencing establishment of CENP-A^Cnp1^ chromatin by sequestering freshly introduced centromeric DNA at SPBs.

### Direct tethering of centromeric DNA to SPBs mediates establishment of CENP-A^Cnp1^ chromatin

Insertion of central domain *cc2* DNA near endogenous centromeres indicates that proximity *in cis* to SPB-centromere clusters enhances CENP-A^Cnp1^ chromatin establishment. If the SPB-centromere cluster creates a nuclear compartment that promotes CENP-A^Cnp1^ assembly then positioning centromeric *cc2* DNA *in trans* near SPBs might also lead to CENP-A^Cnp1^ and kinetochore assembly. To directly test if the SPB-centromere compartment influences CENP-A^Cnp1^ chromatin establishment on centromeric DNA we artificially tethered episomal minichromosomes to SPBs. The inner nuclear membrane (INM) protein Lem2 localizes around the nuclear envelope and also exhibits strong colocalization with SPBs (Figure 5A) (Ebrahimi et al., 2018). Lem2 is also specifically enriched across the central domain of fission yeast centromeres (Barrales et al., 2016; Iglesias et al., 2020). Arrays of *lacO* sites (2.8 kb; ∼90 *lacO* sites) were placed in pcc2, generating pcc2-lacO (Table S2) which was then transformed into cells constitutively expressing a LacI-GFP fusion protein (binds pcc2-lacO) and Lem2 fused to both GFP-binding protein (GBP) and mCherry (Lem2-GBP-mCherry; Figure 5A, B). Therefore, cells expressing both Lem2-GBP-mCherry and LacI-GFP should tether pcc2-lacO to SPBs. Indeed, Lem2-mediated tethering resulted in the pcc2-lacO FISH signal being in close proximity to SPBs in 77% of cells, whereas in the absence of tethering components it was located away from SPBs in >77% of cells examined (Figure 5C, D). Crucially, this Lem2-mediated tethering of pcc2-LacO near SPBs resulted in CENP-A^Cnp1^ incorporation at *cc2* on SPB-adjacent pcc2-lacO, whereas CENP-A^Cnp1^ was not detected on untethered pcc2-lacO or pcc2 itself (Figure 5E). In addition to CENP-A^Cnp1^, the inner kinetochore protein CENP-C^Cnp3^ and outer kinetochore protein Knl1^Spc7^ were also assembled on the *cc2* central domain of pcc2-lacO, but only when it was tethered at SPBs (Figure 5F, G).

**Figure 5.**
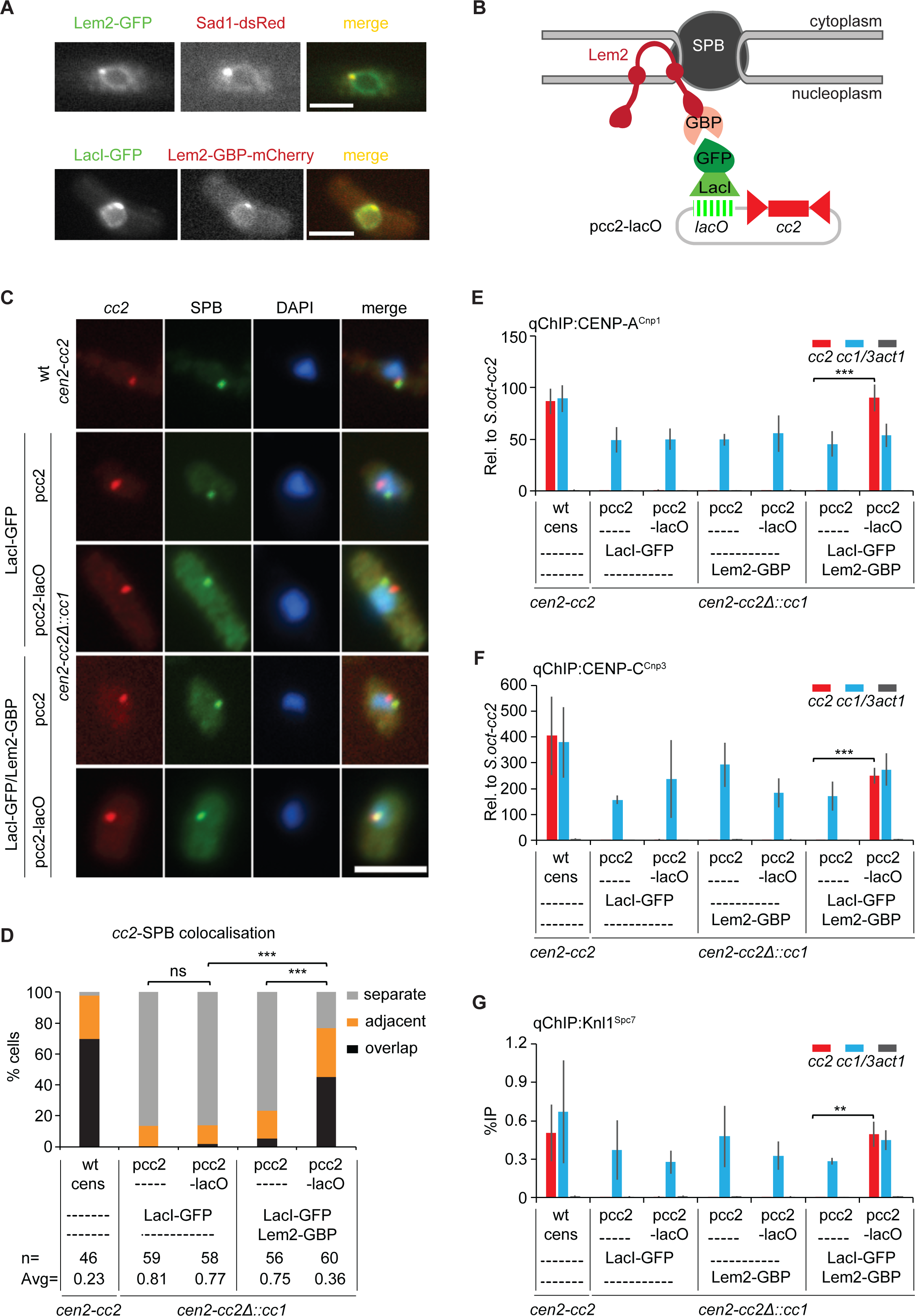
Tethering *cc2* DNA to Lem2 allows CENP-A^Cnp1^ incorporation and kinetochore protein recruitment. (A) Representative images of live cells expressing Lem2-GFP and Sad1-dsRed or LacI-GFP and Lem2-GBP-mCherry. Images were scaled as in Figure 2. Scale bar, 5 μm. (B) Schematic representation of the tethering system used to force pcc2-lacO association with Lem2-GBP-mCherry at the NE and SPB. pcc2-lacO is bound by LacI-GFP and ultimately tethered to Lem2-GBP-mCherry via GFP/GBP interaction. (C) Representative images of *cc2* DNA FISH (red), SPB location (green; anti-Cdc11) and DNA staining (blue, DAPI) in wt cens strain carrying endogenous *cen2-cc2* or *cen2-cc2*Δ::*cc1* strain expressing LacI-GFP or both LacI-GFP and Lem2-GBP-mCherry transformed with pcc2 or pcc2-lacO. Fluorescence of LacI-GFP and Lem2-GBP-mCherry was dissipated by the Immunofluorescence/DNA-FISH procedure and did not contribute punctate signal. Images were scaled as in Figure 1. Scale bar, 5 μm. (D) 3D distances between *cc2* and SPBs (Cdc11), percentage of interphase cells (n, number analyzed) in each category, classified as in Figure 1. Avg, average distance. ns, no significance; ***, p < 0.0001 (Mann-Whitney U test). (E-G) qChIP analyses for CENP-A^Cnp1^ (E), CENP-C^Cnp3^ (F), Knl1^Spc7^ (G) levels at *cc2*, *cc1/3* and *act1* in wt cens strain carrying endogenous *cen2-cc2* or *cen2-cc2*Δ::*cc1* strain expressing LacI-GFP or Lem2-GBP-mCherry or both of them transformed with pcc2 or pcc2-lacO. %IP levels in *S. pombe* were normalized to %IP of *S. octosporus* central core from spiked-in chromatin in (E, F). qChIP results in (G) reported as %IP. Data are mean ± SD (n=3). **, p<0.005; ***, p<0.0005 (Unpaired t-test) (See also Figure S6, S7).

These analyses demonstrate that direct tethering of *cc2* DNA to SPBs enables CENP-A^Cnp1^ chromatin to be established without the need for adjacent heterochromatin. However, Lem2 is not an SPB-specific protein and thus Lem2-mediated pcc2-lacO tethering does not rule out the possibility that the non-SPB fraction of Lem2, localized around the nuclear envelope (Figure 5A), somehow contributes to CENP-A^Cnp1^ and kinetochore protein enrichment. The Alp4 and Alp6 proteins are components of the SPB associated γ-tubulin complex and a proportion of both proteins localize on the nucleoplasmic side of SPBs in interphase (Bestul et al., 2017). Cells expressing Alp4-GBP-mCherry or Alp6-GBP-mCherry fusion proteins and LacI-GFP, transformed with pcc2-LacO were therefore generated (Figure S6A, B). Both Alp4-GBP- or Alp6-GBP-mediated tethering resulted in pcc2-lacO being located close to SPBs in 82-90% of cells, whereas in >79% of cells lacking tethering components pcc2-lacO was located distant from SPBs (Figure S6C-E). Importantly, SPB tethering via Alp4-GBP or Alp6-GBP resulted in CENP-A^Cnp1^ incorporation on the *cc2* region of pcc2-lacO (Figure S6F, G). Thus, the direct tethering of *cc2* DNA to SPBs via SPB-specific components enables CENP-A^Cnp1^ chromatin establishment. The establishment of CENP-A^Cnp1^ chromatin on pcc2-lacO transformed into Lem2-GBP/LacI-GFP-expressing cells was unaffected by the absence of Clr4 H3K9 methyltransferase (*clr4*Δ; Figure S7).

Together these manipulations reveal that in the absence of adjacent heterochromatin the forced localization of centromeric central domain DNA, the native substrate for fission yeast CENP-A^Cnp1^ assembly, to SPBs is sufficient to trigger CENP-A^Cnp1^ chromatin and kinetochore assembly.

### Loss of centromere-SPB association prevents CENP-A^Cnp1^ chromatin establishment

If the *de novo* establishment of CENP-A^Cnp1^ chromatin on centromeric DNA tethered near SPBs depends on the surrounding nuclear compartment, then loss of centromere-SPB association would be expected to hinder CENP-A^Cnp1^ incorporation. The accumulation of Lem2 at SPBs requires the Csi1 protein; in cells lacking Csi1 (*csi1*Δ) Lem2 is mainly localized around the nuclear periphery (Figure 6A) (Ebrahimi et al., 2018). We therefore used *csi1*Δ cells to test if loss of the SPB-associated Lem2 pool affects Lem2-mediated tethering of pcc2-lacO at SPBs (Figure 6A, B). Indeed, pcc2-lacO was located near SPBs in only 23% of *csi1*Δ cells compared to 77% of wild-type cells expressing Lem2-GBP-mCherry and LacI-GFP (Figure 6D, E). Furthermore, *csi1*Δ cells were unable to establish CENP-A^Cnp1^ chromatin on Lem2-tethered pcc2-lacO (Figure 6F). However, CENP-A^Cnp1^ can assemble *de novo* on cc2 of pHcc2 transformed into *csi1*Δ cells (Figure 6G), indicating that Csi1 itself is not required for CENP-A^Cnp1^ establishment. Thus, Lem2 needs to be concentrated at SPBs in order to induce CENP-A^Cnp1^ incorporation on tethered centromeric DNA.

**Figure 6.**
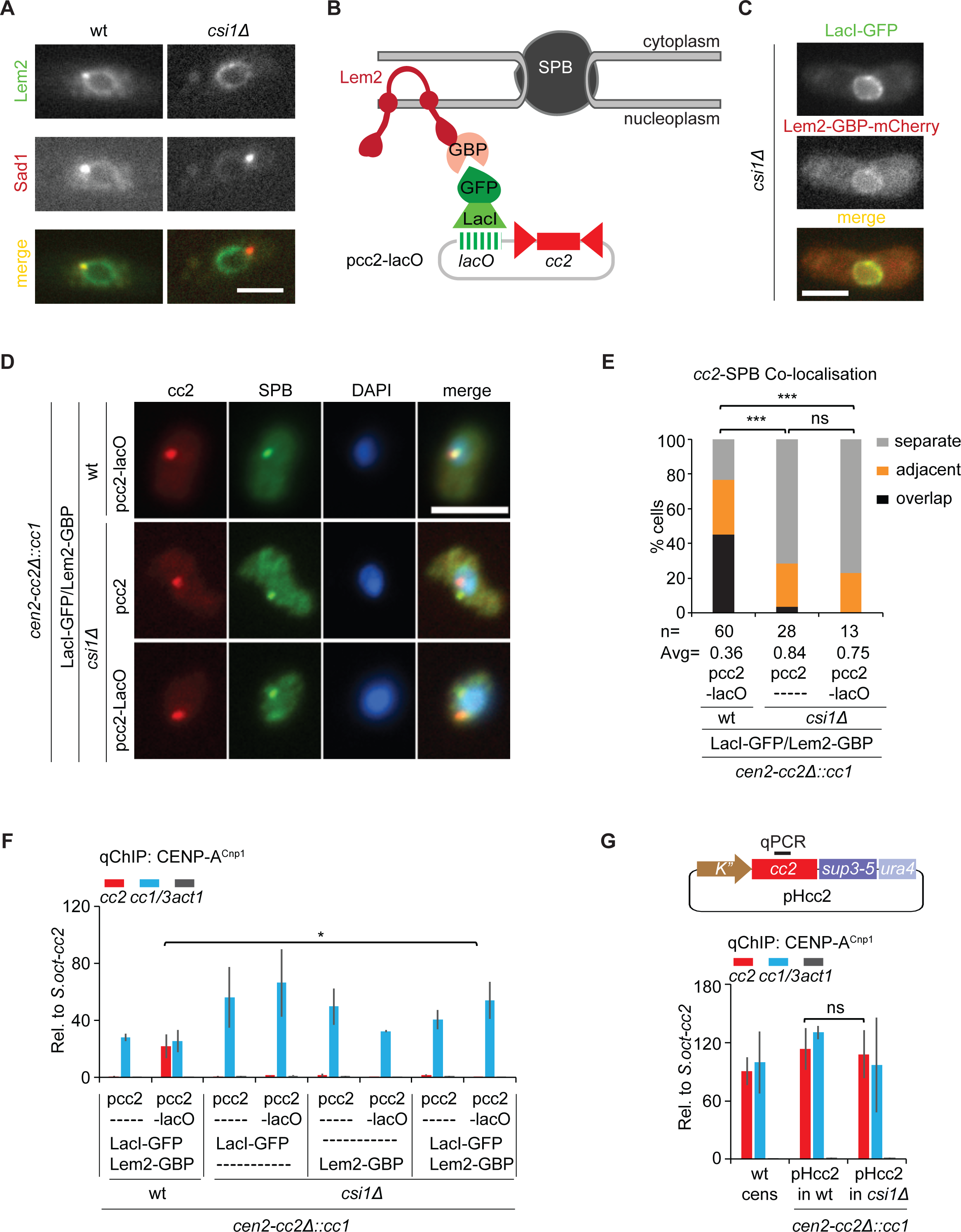
Loss of Csi1 prevents CENP-A^Cnp1^ chromatin establishment on Lem2-tethered pcc2-*lacO*. (A, C) Representative images of live wt and *csi1*Δ cells expressing Lem2-GFP and Sad1-dsRed (A) or LacI-GFP and Lem2-GBP-mCherry (C). Images were scaled as in Figure 2. Scale bar, 5 μm. (B) Forced association of pcc2-lacO with Lem2-GBP-mCherry at NE in *csi1*Δ using same tethering system as in Figure 5. In *csi1*Δ, pcc2-lacO is expected to detach from the SPB due to loss of Lem2 from SPB. (D) Representative images of *cc2* DNA FISH (red), SPB location (green; anti-Cdc11) and DNA staining (blue, DAPI) wt or *csi1*Δ strains expressing both LacI-GFP and Lem2-GBP-mCherry transformed with pcc2 or pcc2-lacO. Images were scaled as in Figure 1. Scale bar, 5 μm. (E) Percentage of interphase cells (n, number analyzed) displaying distinct degrees of *cc2* DNA colocalization with SPBs (Cdc11). Cells were classified into three groups as in Figure 1. Avg, average distance. ns, no significance; ***, p < 0.0001 (Mann-Whitney U test). (F, G) qChIP analyses for CENP-A^Cnp1^ at *cc2*, *cc1/3* and *act1* in indicated strains transform with pcc2 or pcc2-lacO (F) or pHcc2 (G). qChIP primer site on pHcc2-borne *cc2* is indicated as black bar above plasmid map (G). %IP levels in *S. pombe* were normalized to %IP of *S. octosporus* central core from spiked-in chromatin. Data are mean ± SD (n=3). ns, no significance; *, p<0.05 (Unpaired t-test).

**Figure 7.**
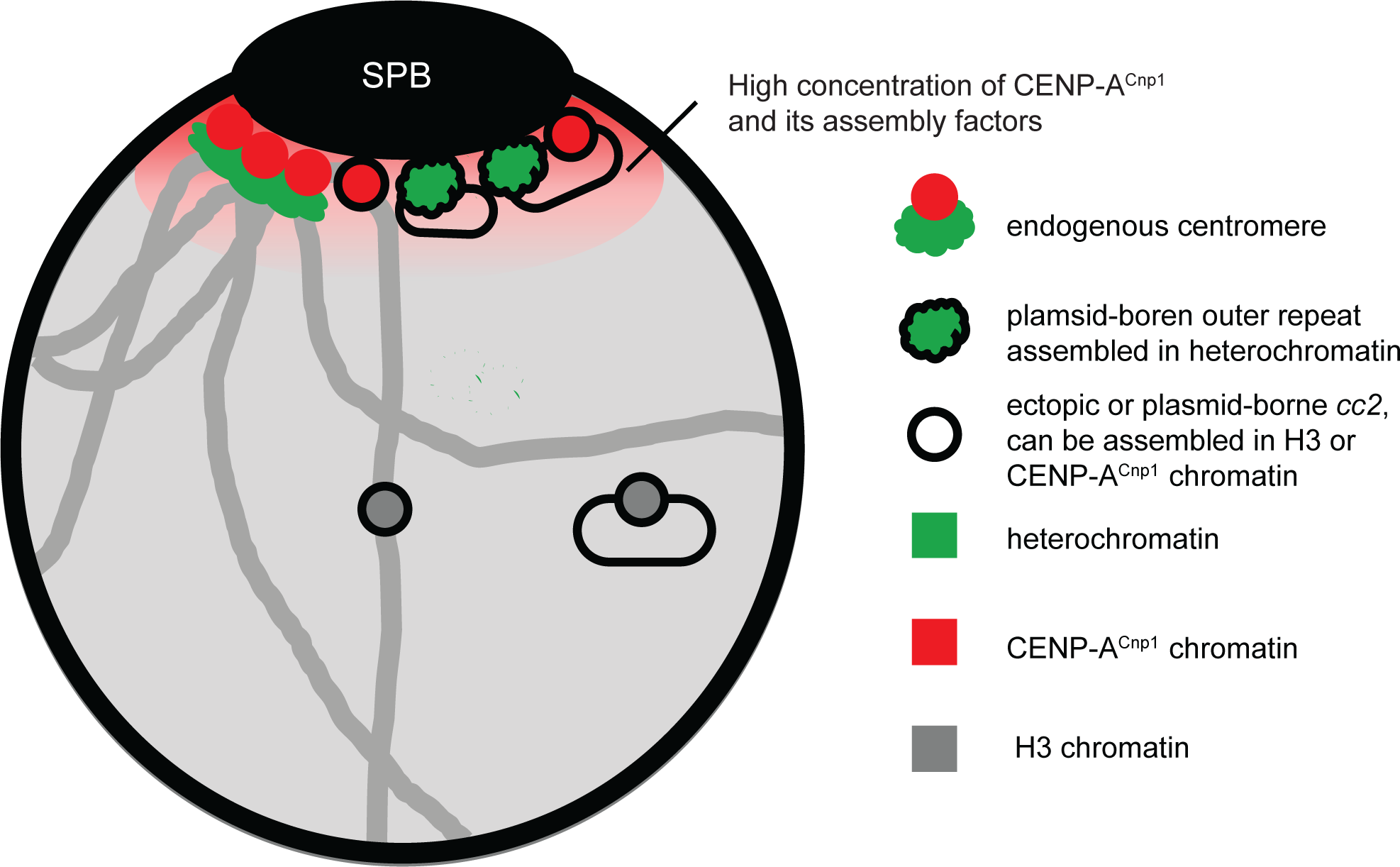
Model: Centromere identity is influenced by nuclear spatial organization. Due to clustering of endogenous centromeres (CENP-A^Cnp1^-assembled central domains; red circles; heterochromatic outer repeats: green) at SPBs and incorporation of CENP-A^Cnp1^ at centromeres in G2, the zone around SPBs forms a nuclear sub-compartment rich in CENP-A^Cnp1^ and its assembly factors (red shaded cloud). Ectopic central domain (outlined circles) inserted at centromere-proximal sites are exposed the high-CENP-A^Cnp1^ SPB/centromere sub-compartment, promoting *de novo* incorporation of CENP-A^Cnp1^, unlike centromere-distal locations. Similarly, only minichromosomes bearing heterochromatin, which mediates association with the SPB, exposes adjacent central domain to the high-CENP-A^Cnp1^ SPB/centromere sub-compartment, resulting in CENP-A^Cnp1^ incorporation. Heterochromatin, green; CENP-A^Cnp1^, red; neutral H3 chromatin, grey.

Together these data indicate that pericentromeric heterochromatin is sufficient to mediate frequent contact with SPBs where centromeres and CENP-A^Cnp1^ assembly factors are concentrated. We conclude that heterochromatin promotes exposure of adjacent *cc2* centromere DNA to this CENP-A^Cnp1^ assembly factor-rich nuclear sub-compartment, thus ensuring the assembly of CENP-A chromatin and kinetochores.

## Discussion

Assembly of CENP-A chromatin is epigenetically regulated. Here we demonstrate that, in addition to the impact of chromatin context and prior CENP-A history, spatial location within the nucleus is an epigenetic influence on the chromatin fate of centromeric DNA. We show that heterochromatin causes minichromosomes to localize near SPBs, providing a likely explanation for the role of heterochromatin in promoting CENP-A^Cnp1^ chromatin establishment on adjacent centromeric sequences. By placing a CENP-A^Cnp1^ assembly-competent sequence (*cc2*) in various spatial contexts we demonstrate that being in the vicinity of centromere clusters at SPBs triggers CENP-A^Cnp1^ chromatin establishment.

Despite epigenetic factors being important in the establishment of CENP-A chromatin, certain sequences are preferred, including human *α*-satellite arrays and fission yeast central domains. Rather than the precise sequence being critical, evidence suggests that innate properties of central domain regions, such as their unusual transcriptional landscape and high rates of histone H3 turnover, are permissive for CENP-A^Cnp1^ incorporation into chromatin (Catania et al., 2015; Shukla et al., 2018; Singh et al., 2020). Although central domain sequences are the preferred substrate for CENP-A^Cnp1^ assembly in fission yeast, *de novo* assembly of CENP-A^Cnp1^ chromatin is context dependent. Outer repeat-directed or synthetic heterochromatin promotes CENP-A^Cnp1^ chromatin establishment on adjacent central domain DNA (Folco et al., 2008; Kagansky et al., 2009). CENP-A^Cnp1^ overexpression induces *de novo* CENP-A^Cnp1^ chromatin establishment on plasmid-based minichromosomes devoid of heterochromatin and carrying only central domain sequences (Catania et al., 2015).

We have previously suggested two models to explain these observations (Catania et al., 2015; Choi et al., 2012; Folco et al., 2008). In the ‘modifier’ model, heterochromatin performs a chromatin-directed role such as recruitment of histone-modifying enzymes or remodelers that influence histone dynamics to favor CENP-A^Cnp1^ incorporation on adjacent central domain regions. In this scenario, CENP-A^Cnp1^ overexpression would shift the equilibrium away from transcription-dependent histone H3 recycling and towards CENP-A^Cnp1^ deposition. In the ‘positioning’ model, the role of heterochromatin, due to its own localization, would place central domain DNA at a nuclear location permissive for CENP-A^Cnp1^ deposition, such as a compartment exhibiting high levels of CENP-A^Cnp1^ and associated assembly factors. In this model, overexpression of CENP-A^Cnp1^ would bypass heterochromatin’s function by making a greater proportion of nuclear space permissive for CENP-A^Cnp1^ assembly.

Here we have utilized FISH to demonstrate that minichromosome-borne heterochromatin preferentially locates close to SPBs. We hypothesize that any sequence positioned at this location will be exposed to high concentrations of CENP-A^Cnp1^ and its assembly factors because centromeres are clustered at SPBs for most of the cell cycle. However, only sequences such as centromeric central domain DNA, with embedded properties that drive transcription-coupled H3 replacement with CENP-A^Cnp1^, actually incorporate CENP-A^Cnp1^ (Shukla et al., 2018).

Support for the hypothesis that heterochromatin’s role in CENP-A^Cnp1^ establishment is to position central domain within the SPB-centromere cluster compartment of the nucleus is provided by our finding that centromeric *cc2* DNA inserted close to endogenous or neo-centromeres assembled CENP-A^Cnp1^ chromatin, whereas *cc2* inserted at locations far away from centromeres did not. The positioning of *lys1* and *itg8* close to SPBs in wild-type and neocentromere-containing cells, respectively, correlates with the incorporation of CENP-A^Cnp1^ on *cc2* when inserted at these sites. Although the failure of CENP-A^Cnp1^ to assemble on centromere-distal sites such as *ade3* could be attributed to selection against deleterious dicentric formation on this endogenous chromosome, we have previously shown that *cc2* present on the arm of a 530-kb non-essential linear minichromosome also does not normally assemble CENP-A^Cnp1^. However, that minichromosome is capable of dicentric formation because overexpressed CENP-A^Cnp1^ incorporates into *cc2* and causes missegregation (Catania et al., 2015). Thus, placing central domain DNA near centromeres *in cis* results in CENP-A^Cnp1^ incorporation. Moreover, direct tethering of minichromosome-borne central domain DNA *in trans* to SPB-associated proteins also triggered *de novo* assembly of CENP-A^Cnp1^ chromatin, bypassing the requirement for heterochromatin. Thus, when susceptible sequences are positioned in the vicinity of SPBs, establishment of CENP-A^Cnp1^ chromatin is uncoupled from the presence of heterochromatin. These observations indicate that nuclear positioning is an epigenetic factor important for establishing centromere function and the function that heterochromatin provides is positioning information.

Fission yeast neocentromeres arise most frequently in subtelomeric regions and immature neocentromeres near rDNA can be stabilized by relocation to subtelomeric regions or upon acquisition of adjacent heterochromatin (Ishii et al., 2008; Ogiyama et al., 2013). When overexpressed, CENP-A^Cnp1^ is incorporated at moderate levels over subtelomeric regions (Castillo et al., 2013). Therefore, subtelomeric regions represent favored, but secondary, sites for CENP-A^Cnp1^ and kinetochore assembly. H3K9me-dependent heterochromatin is normally assembled adjacent to telomeres (Kanoh et al., 2005). During interphase fission yeast telomeres are attached to the nuclear envelope via inner nuclear membrane proteins Bqt3 and Bqt4 (Chikashige et al., 2009). Moreover, Hi-C analysis detects frequent contacts between telomere and centromere regions (Mizuguchi et al., 2014). We suggest that as consequence of their association with the nuclear periphery, subtelomeric regions perform a highly constrained exploration of the nucleus, making them more likely to meet the SPB-centromere cluster which exposes them to the immediate nuclear compartment containing high levels of CENP-A^Cnp1^ assembly factors. Thus, association with the nuclear envelope offers an attractive explanation for the subtelomeric location of most fission yeast neocentromeres. Neocentromeres arise at telomeres even in the absence telomeric heterochromatin (Ishii et al., 2008), suggesting that it is telomere anchoring at the nuclear envelope, not subtelomeric heterochromatin, that designates them as secondary CENP-A^Cnp1^ assembly sites.

In fission yeast, centromeres are clustered at the SPB throughout the cell cycle, apart from during mitosis, returning to the SPB in anaphase. CENP-A^Cnp1^ and several CENP-A^Cnp1^ assembly factors and chaperones such as Scm3^HJURP^, Mis16^RbAP46/48^, Mis18, Eic1/Mis19 are concentrated on centromeres around the SPB during interphase (Hayashi et al., 2004; Pidoux et al., 2009; Subramanian et al., 2014; Williams et al., 2009). Mammalian centromeres are not localized close to centrosomes (SPB equivalent) during most of the cell cycle. However, after mitotic chromosome segregation, mammalian centromeres transiently cluster at spindle poles in late anaphase/telophase, subsequently dispersing during G1 (Gerlich et al., 2003). Centromere clustering is also pronounced in plants that exhibit an overt ‘Rabl’ configuration where centromeres and telomeres are clustered at opposite sides of interphase nuclei (Oko et al., 2020). Intriguingly, the Mis18 CENP-A assembly complex is normally recruited to human centromeres in late anaphase/telophase prior to arrival of the HJURP CENP-A chaperone and new CENP-A incorporation in early G1 (Zasadzińska and Foltz, 2017), although loss of CDK (cyclin-dependent kinase) regulation allows premature CENP-A deposition in G2 cells (Stankovic et al., 2017). Therefore, the centromeres of complex eukaryotes are briefly clustered together at precisely the time when assembly factors are recruited to centromeres. This spatio-temporal co-ordination may maximize the local concentration of CENP-A and its assembly factors to ensure the efficient removal of H3 placeholder nucleosomes and replenishment of CENP-A nucleosomes in centromeric chromatin (Dunleavy et al., 2011).

Once CENP-A^Cnp1^ chromatin and kinetochores are assembled at fission yeast centromeres, it is clear that heterochromatin-independent connections with SPBs are established. Centromeres remain clustered at SPBs in the absence of pericentromeric H3K9me-dependent heterochromatin but SPB-centromere clustering is disrupted when essential kinetochore components such as Mis6 or Nuf2 are defective (Appelgren et al., 2003; Saitoh et al., 1997). Thus, once assembled, an intact interphase kinetochore structure rather than pericentromeric heterochromatin appears to provide the main physical link between functional centromeres and SPBs.

Here we have demonstrated that the specific location of centromere sequences within nuclei (i.e. their spatial context) exerts an epigenetic influence on the eventual CENP-A chromatin state attained by specific DNA sequences. Our analyses demonstrate that the SPB-centromere cluster forms a sub-compartment within the nucleus that promotes CENP-A and kinetochore assembly on DNA sequences presenting the required features to facilitate CENP-A chromatin assembly in place of canonical H3 chromatin. Thus, spatial positioning in the nucleus is a hitherto unrecognized epigenetic determinant of centromere identity.

## Author Contributions

Conceptualization, W.W., A.L.P. and R.C.A.; Methodology, W.W., T.M., D.A. and A.L.P.; Investigation, W.W.; Visualization, W.W.; Writing – First Draft, W.W.; Writing – Review & Editing, W.W., A.L.P. and R.C.A.; Funding Acquisition, R.C.A.; Supervision, A.L.P. and R.C.A.

## Acknowledgments

We are grateful to members of the Allshire lab especially Manu Shukla, Nitobe London and Dominik Hoelper for helpful suggestions and comments on the manuscript and Sharon White for organizational support. Takeshi Urano is thanked for the 5.1.1 (H3K9me) antibody; Ken Sawin for the Cdc11 antibody; Kevin Hardwick for the Spc7 antibody; Julie Cooper for the plasmid pF6a-GBP-mCherry-Hyg; and Nick Rhind for *S. octosporus* strain. W.W. is supported by the Darwin Trust of Edinburgh. R.C.A. is a Wellcome Principal Research Fellow (095021; 200885); the Wellcome Centre for Cell Biology is supported by core funding from Wellcome (203149). The Allshire lab dedicates this study to the memory of our dear colleague and friend Sasha Kagansky whose research and insights were an inspiration for this project.

## Methods

### Yeast strains

Yeast strains used in this study and their genotypes are listed in Table S1.

Standard genetic and molecular methods were used as described (Moreno et al., 1991). All ectopic *cc2* insertions were made in *cc2*Δ::*cc1* strains (Catania et al., 2015) by integrating linear *cen2* central domain constructs (∼880 bp *imr2L,* -6.8 kb *cc2* and ∼920 bp *imr2R*, abbreviated as *cc2*) by homologous recombination (HR). pMC52 (Table S2), bearing 8.6 kb of *cc2* and kanMX6 selection cassette, was used as a starting plasmid for linear *cc2* constructs. Two flanking DNA fragments of the desired target locus for *cc2* insertions were amplified using primers listed in Table S3 by PCR. Restriction enzyme *Kpn*I/*Xho*I-digested first fragment was cloned into *Kpn*I/*Xho*I-digested pMC52, which were then digested by *Sac*I/*Msc*I and ligated with *Sac*I/*Msc*I-digested second PCR fragment by T4 DNA ligase (M0202S; NEB). Linear *cc2* constructs were obtained by *Sac*I/*Kpn*I digestion of the resulting plasmids and transformed into desired strain for *cc2* insertion.

For the construction of Lem2/Alp4/Alp6-GBP-mCherry and Lem2-GFP, the GBP-mCherry-hygMX6 and GFP-natMX6 cassette in plasmid pFA6a-GBP-mCherry-hygMX6 (Fernández-Álvarez et al., 2016) and pFA6a-GFP-NatMX6 were amplified by PCR and integrated into genome by HR (Bähler et al., 1998).

*clr4*Δ mutant was created by CRISPR/Cas9 method as described previously (Torres-Garcia et al., 2020). Briefly, *clr4* gene specific sgRNA was cloned into Cas9 containing pLSB-KAN plasmid by Golden Gate Assembly kit (E1601S, NEB). The resulting plasmid *clr4*-pLSB-KAN and *clr4* HR template obtained by annealing primer pair WW748-clr4-HR-F/WW749-clr4-HR-R (Table S3) were co-transformed into *S. pombe* by sorbitol-electroporation method.

Transformants were grown on appropriate selection plates and screened for correct integration or *clr4*Δ mutant by yeast colony PCR using primers listed in Table S3. All plasmids and primers used in this study are listed in Table S2, Table S3 respectively.

### Yeast growth medium and conditions

All strains were grown at 32°C in YES (Yeast Extract with Supplements) rich medium or PMG (Pombe Minimal Glutamate) minimal medium, as appropriate. Selection for dominant markers was performed on YES medium supplemented with 100 μg/ml clonNAT (96736-11-7, Werner BioAgents), 100 μg/ml G418 (10131027, Gibco), or 123 μg/ml HygMX6 (31282-04-9, Duchefa Biochemie). *clr4*Δ transformants were selected on YES supplemented with G418 plate and re-streaked to non-selective YES medium to allow loss of plasmid clr4-pLSB-KAN. Transformants with *cc2* insertions were selected on YES supplemented G418. Plasmids pcc2 (pMC2; carrying 8.6 kb of *cc2*) and pcc2-LacO (pMC12; carrying 8.6 kb of *cc2* and 2.8 kb of *lacO*) were selected on YES containing 100 μg/ml G418 in wt strains or on PMG-uracil in *csi1*Δ (*csi1*Δ::*ura4*) strain. Strains carrying plasmid pHet (pMC183; carrying 2 kb of *K”* repeats) or pHcc2 (H denotes 5.6 kb of *K”* repeats, *cc2* denotes 8.6 kb of *cc2*) were selected on YES supplemented with clonNAT or PMG-adenine-uracil medium respectively.

### Bacteria

DH5α E. coli strains (C2987H, NEB) were grown in LB medium at 37°C. E. coli competent cells carrying plasmids were selected on LB agar plates supplemented with 100 μg/ml of ampicillin or LB liquid supplemented with 50 μg/ml Carbenicillin (10177012; Invitrogen).

## Methods details

### Yeast genetic crosses

To obtain desired genotypes, two strains with opposite mating type (h+/h-) were mixed and grown on the nitrogen starved ME plate for sporulation at 32°C for 2 days. Asci were digested in glusulase (NEE-154, NEN) to release spores that were then plated on appropriate selective medium and grown at 32°C.

### Yeast colony PCR

Yeast strains were suspended in SPZ buffer (1.2 M sorbitol, 100 mM sodium phosphate and 2.5 mg/ml Zymolyase-100T (08320932, MP Biomedicals)) and incubated at 37°C for 30 min. The resulted mixtures were used as PCR template for strain genotyping by Roche FastStart™ Taq polymerase PCR kit (12032953001, Roche) supplemented with primers.

### Yeast transformation

Yeast cells were transformed by sorbitol-electroporation method. Log phase cultures were harvested and resuspended in pre-transformation buffer (25 mM DTT, 0.6 M sorbitol and 20 mM HEPES, pH7.6) and incubated at 32°C with 180 rpm shaking for 10 min. Cells were washed three times in ice-cold 1.2 M sorbitol, mixed in an ice-cold cuvette with 200 ng of plasmid DNA or purified DNA fragments obtained by QIAquick PCR Purification Kit (28104, QIAGEN) and then pulsed by an electroporator (Bio-Rad Gene Pulser II) at a setting of 2.25kV, 200Ω and 25μF. Cells were either directly plated on medium with prototrophic selection directly or grown overnight in non-selective liquid before selection for antibiotic resistance (G418/cloNAT/HygMX6). Single colonies were isolated from selective medium.

### Minichromosome establishment assay

Medium-sized colonies carrying circular plasmid-based minichromosome pHcc2 were replica-plated from PMG-adenine-uracil to PMG-low-adenine (10 μg/ml adenine) and incubated at 32°C for 2 days to determine functional centromere establishment frequency. Plasmid pHcc2 contains *sup3e* tRNA selection marker that suppresses *ade6-704* mutation within strains, thus colony color on these PMG-low-adenine plates will indicate minichromosome loss (red colonies) or retention (white/pale pink colonies). In the absence of centromere establishment, minichromosomes behave as episomes that are rapidly lost. Minichromosomes that have established functional centromere segregate efficiently during mitosis. Minichromosomes occasionally integrate at genome will give a false-positive white phenotype. To assess the frequency of such integration events and to confirm establishment of centromere segregation function, colonies giving the white/pale-pink phenotype upon replica plating were re-streaked to single colonies on PMG-low-adenine plates. Red/white sectored colonies are indicative of centromere function with low levels of minichromosome loss, whereas pure white colonies are indicative of integration into endogenous chromosomes. Therefore, the percentage of sectored colonies number over the total colony numbers was used to represent centromere establishment frequency on minichromosome pHcc2.

### Quantitative chromatin immunoprecipitation (qChIP)

Three independent cell cultures were grown in appropriate medium until log phase and fixed in 1% formaldehyde (F8775, MERCK) for 15 min followed by quenching in 125 mM Glycine (G8790, MERCK) at room temperature. ChIP was performed as previously described (Castillo et al., 2007). 2.5x10^8^ cells were used for each ChIP. Briefly, cells were lysed by bead beating (Biospec) in 350 μl Lysis Buffer (50 mM Hepes-KOH pH 7.5, 140 mM NaCl, 1 mM EDTA, 1% (v/v) Triton X-100 and 0.1% (w/v) sodium deoxycholate) supplemented with 3.5 μl of 100 mM PMSF (329-98-6, MERCK) and 3.5 μl of 100 mM yeast protease inhibitor (P8215, MERCK). Where indicated, ∼5x10^7^ fixed, lysed *S. octosporus* cells (Rhind et al., 2011) were added to each initial crude cell lysates as a spike-in control. Crude cell lysates were sonicated using a Bioruptor (Diagenode) at 4°C on high voltage for 20 min (20 cycles of 30 s ON/OFF), followed by centrifugation at 13000 rpm for 10 min to pellet cell debris. The resulting supernatant was used for following steps.

For H3K9me2 ChIP, 10 μl lysate was retained as crude ‘input’ sample, whereas 300 μl of the remaining lysates were incubated overnight with 20 μl of washed protein G Dynabeads (10009D, Thermo Fisher Scientific) and 1 μl of mouse anti-H3K9me2 (mAb5.1.1, gift from Takeshi Urano).

For CENP-A^Cnp1^/CENP-C^Cnp3^/Knl^Spc7^ ChIP, lysates were precleared for 1 h with 25 μl of washed protein-G agarose beads (11243233001, Roche) and 10 μl of precleared lysate was retained as crude ‘input’ sample. 300 μl of the remaining pre-cleared lysates were incubated overnight with appropriate amount of antibody (10 μl of sheep CENP-A^Cnp1^, CENP-C^Cnp3^, CENP-K^Sim4^ serum (Catania et al., 2015) (in-house preparation), 3 μl of affinity-purified sheep anti-Spc7 (a gift from Kevin Hardwick) and 25 μl of protein-G agarose beads.

After immunoprecipitation, the crude “IP” samples on beads were washed in Lysis Buffer, Lysis Buffer supplemented with 500 mM NaCl, Wash Buffer (10 mM Tris-HCl pH 8, 250 mM LiCl, 0.5% IGEPAL NP40 (56741, MERCK) 0.5% (w/v) sodium deoxycholate and 1 mM EDTA) and TE Buffer (10mM Tris-HCl pH 8, 1 mM EDTA). DNA was recovered from input and IP samples using Chelex resin (1421253, BioRad). Quantitative PCR reactions (qPCR) were performed using a LightCycler 480 SybrGreen Master Mix (04887352001, Roche) and analyzed using Roche LightCycler software (version 1.5.1.62). Primers used for qPCR are listed in Table S3. ChIP enrichments on regions of interest were calculated as the ratio of “IP” sample to the corresponding “input” sample using the ΔCT method and represented as %IP. Where indicated, for spike-in qChIPs, %IP levels in *S. pombe* were normalized to %IP from spiked-in *S. octosporus* chromatin (specified in the figure legends).

### Fluorescence microscopy

Live fission yeast cells were mounted on a 2% agarose pad formed on 1 mm SuperFrost slides (Thermo Scientific) whereas fixed cells (immunofluorescence and DNA FISH) were mounted in VECTASHIELD Mounting Medium (H-1000-10, Vector Laboratories) on 1 mm Polysine slides (Thermo Scientific). Microscopy was performed with Nikon Ti2 inverted microscope equipped with a ×100 1.49 NA CFI Plan Apochromat TIRF objective, Lumencor Spectra X light source (Lumencor, Beaverton, OR USA) and a Photometrics Prime 95B camera (Teledyne Photometrics, Birmingham, UK), all controlled by Nikon NIS Elements software version 5.21.03 (RRID:SCR_014329). Filter sets from Semrock (Semrock, Rochester, New York, USA) were used to image Lem2-GFP, LacI-GFP, Alexa Fluor 488 (A-11015, Invitrogen) at excitation 488 nm, emission 535 nm, Sad1-dsRed, Rhodamine at excitation 554 nm, emission 590 nm, Lem2/Alp4/Alp6-mCherry at excitation 578 nm, emission 630 nm and DAPI, excitation 378 nm, emission 460 nm. A Mad City nano drive (Mad City Labs, Madison, WI, USA) was used to produce whole cell 3 dimensional (3D) images with a step size of 0.3 μm. All images were processed by Fiji software (RRID:SCR_002285). Live cell images were scaled relative to the maximum intensity in the set of images to allow comparison between images, but fixed cell images were scaled relative to the maximum value of histogram (specified in figure legends).

### Immunofluorescence/DNA FISH

For Immunofluorescence/DNA FISH, cells were initially subjected to a similar Immunofluorescence protocol as described previously with some modifications (Castillo et al., 2007) and subsequent FISH process. Briefly, log phase yeast cultures were fixed with 3.7% formaldehyde for 7 min at room temperature, washed by PEM buffer (100 mM PIPES pH 7, 1 mM EDTA, 1 mM MgCl_2_) and PEMS buffer (100 mM PIPES pH 7, 1 mM EDTA, 1 mM MgCl_2_, 1.2 M Sorbitol), followed by cell-wall digestion in PEMS buffer supplemented with 1 mg/ml Zymolyase-100T and 1 mg/ml Lallzyme (Lallzyme-MMX, Litmus Wines) at 37°C for 90 min. After permeabilization in PEMS containing 1% Triton X-100 for 5 min at room temperature, cells were washed, blocked in PEMBAL (PEM containing 1% BSA (A0281, MERCK), 0.1% sodium azide, 100 mM lysine hydrochloride (657-27-2, MERCK)) for 1 h. Cells were then incubated overnight at 4 °C with 1:500 anti-Cdc11 (Castillo et al., 2007) (a SPB protein; gift from Ken Sawin) or 1:500 anti-CENP-A^Cnp1^ (in-house preparation) in 500 μl of PEMBAL. Cells were then washed three times with PEMBAL and incubated overnight with 1:500 Alexa-488-coupled donkey anti-sheep secondary antibody (A-11015, Invitrogen) in 500 μl of PEMBAL. Cells were then washed in PEMBAL and PEM buffer and re-fixed in 3.7% formaldehyde and 0.25% glutaraldehyde (111-30-8, MERCK) for 15 min, washed with PEM buffer and treated with 1 mg/ml sodium borohydride in PEM buffer. After incubation with 2 μl of 10 mg/ml RNase A (19101, Qiagen) in 100 μl of PEMBAL at 37 °C for 2h, cells were denatured in 100 μl of freshly prepared 0.1 M NaOH for 1 min and hybridized with 2 μl of DNA FISH probe in 100 μl hybridization buffer (10% Dextran sulphate (D8906, MERCK), 50% deionized formamide (S4117, MERCK), 2XSSC, 5X Denhardts (D2532, MERCK), 0.5 mg/ml denatured salmon sperm DNA) at 37 °C overnight.

For *lys1*, *itg8* and *neo1R* FISH probe, a ∼12.5 kb region (ChrI: 3,727,604-3,737,389 and ChrI: 3,739,857-3,742,327) spanning *lys1* gene, ∼12.5 kb region (ChrI: 5,495,975-5,508,459) spanning *itg8* locus (ChrI: 5,500,986-5,502,881) and ∼16.3 kb region (ChrI: 5,513,871-5,530,124) within *neo1R* CENP-A^Cnp1^ domain were amplified by PCR using primers listed in Table S3 respectively. Plasmid pMC52, pMC1 was used to make *cc2* and plasmid backbone DNA FISH probes, respectively. *cc2* DNA FISH probe was used to locate *cc2* at endogenous *cen2*, *lys1* and plasmid pcc2 and pHcc2, while plasmid backbone probe was used to locate pHet. FISH probes were obtained by DIG labeling 500 ng DNA (PCR products or plasmids) using DIG-Nick Translation Mix (11745816910, Roche) supplemented with 1 μl of 1:50 diluted DNase I (AM2222, Ambion).

After hybridization with DNA FISH probe, cells were washed with 2XSSC containing 0.1% sodium azide and incubated with 1:100 sheep anti-DIG-Rhodamine (11207750910; Roche) in 100 μl of PBS-BAG (PBS buffer supplemented with 1% BSA (A0281, MERCK), 0.1% sodium azide and 0.5% cold water fish gelatin (G7765, MERCK)) at room temperature overnight. Cells were finally stained with 4’,6-diamidino-2-phenylindole (DAPI), mounted in VECTASHIELD Mounting Medium on Polylysine slides and imaged using Nikon NIS Elements software (version 5.21.03) on a Nikon Ti2 inverted microscope as indicated above. All images are scaled relative to the maximum value of histogram.

### 3D distance measurements

3D distances between spots in two channels (green and red): Cdc11/CENP-A^Cnp1^ (green) and DNA FISH (red) or *lys1:lacO* (*ade3:lacO*)/LacI-GFP (green) and Sad1-dsred (red), were measured by Fiji using in-house script (https://doi.org/10.5281/zenodo.5657360). Briefly, the center of spot in each channel were determined in X-Y using the Fiji “Find Maxima…” function with same threshold (Prominence>500), applied to a Z-projection. The Z-positions of each spot were then determined as the slice with the maximum pixel intensity at each X-Y position. The distance to the nearest red spot for each green spot was reported if within 3 µm representing the diameter of the fission yeast nucleus. If no red spot was detected with 3 µm then that green spot was not included in the analysis. Distances between the resulting spots in each channel were measured by equation:

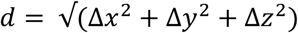

Live mono-nuclear cells 8-12 μm in length and only one SPB (Sad1-dsRed) nucleus-associated dot were recognized as G2 cells and subjected to distance measurements between LacI-GFP (binds to *lys1:lacO* or *ade3:lacO*) and Sad1-dsRed. For immunofluorescence/DNA FISH, mononuclear cells with nuclear green-red spot pairs and only one SPB (Cdc11) or centromere cluster (CENP-A^Cnp1^) spot were recognized as interphase cells and retained for distance measurement between DNA FISH locus (red) and protein Cdc11 or CENP-A^Cnp1^ (green).

### Quantification and Statistical Analysis

All quantification and statistical details of experiments are described in the figure legends or in the methods section. The qChIP results are obtained from more than 3 independent experimental replicates (n*≥*3) and represented as mean ± SD (standard deviation, error bars). Significance of the differences in qChIP results was evaluated using Unpaired t-test with a p value threshold < 0.05, by Prism Version 9.1.0 software (RRID:SCR_002798). 3D distance measurement results were obtained by analyzing n number of interphase cells from 3 independent experimental replicates. Average distance for each strain were calculated and indicated as “Avg” (specified in figure legends). Cells were classified into three groups according to the distance: overlap (≤0.3 μm), adjacent (0.3-0.5 μm) or separate (0.5-3 μm). The results were reported as percentage of cells (% cells) in each group. For statistical significance analysis of distance data, Mann-Whitney U test with a p value threshold <0.01was performed by Prism Version 9.1.0 software (RRID:SCR_002798) and the detailed results were showed in Table S4.

## Supplemental Text and Figures

**Figure S1.**
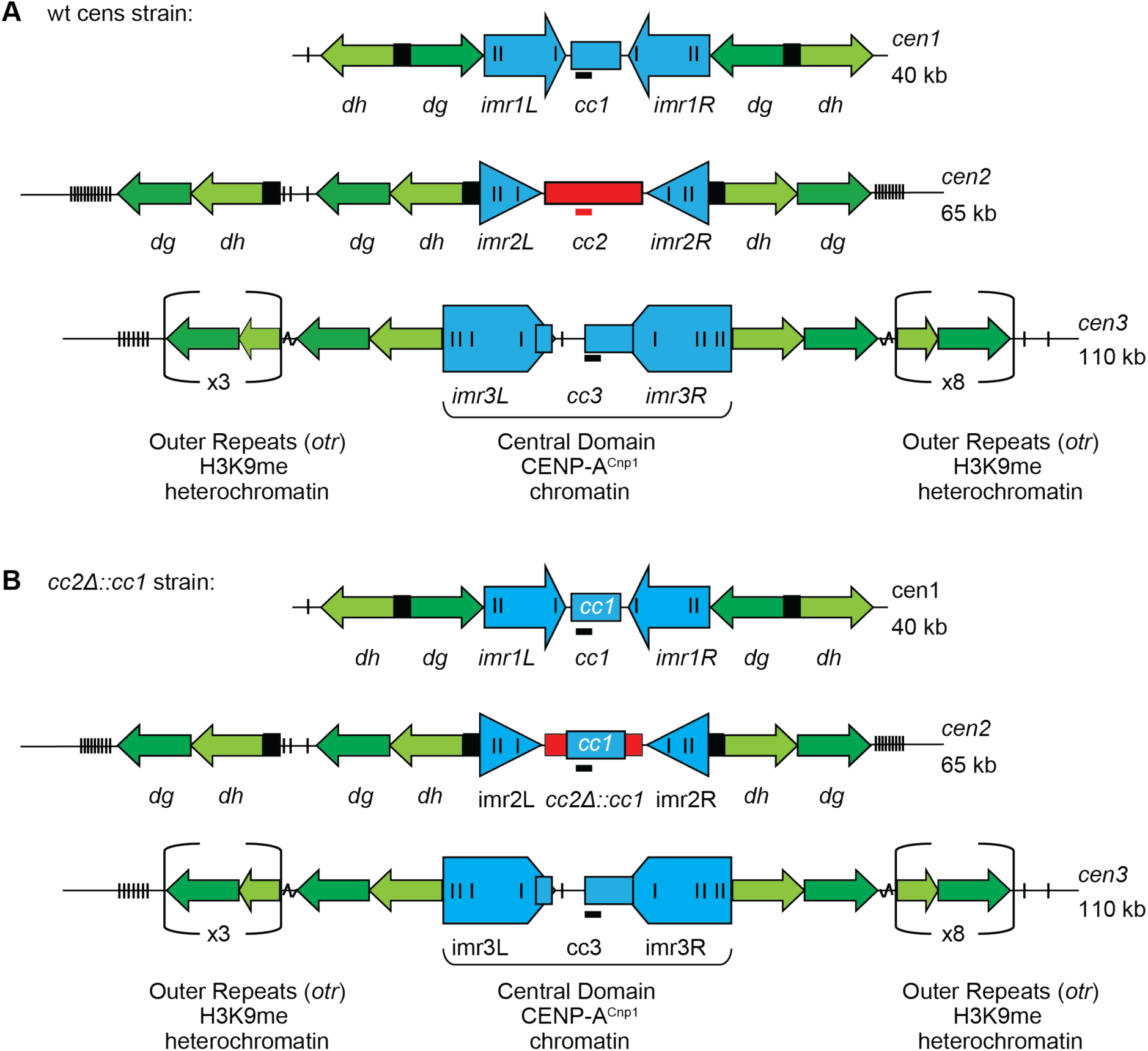
Domain organization of centromeres in fission yeast. Relates to Figures 1-7. (A, B) Schematic representation of the three endogenous centromeres in wt-cens (A) or *cc2*Δ::*cc1* strains (B). Each centromere consists of two distinct domains: a central domain assembled in CENP-A^Cnp1^ chromatin harbouring a central core (*cc*) DNA and flanking innermost repeats (*imr*), which are surrounded by various repetitive DNA elements known as the outer repeats (*otr-dg/dh*) assembled in H3K9me-dependent heterochromatin (Allshire and Ekwall, 2015). In *cc2*Δ::*cc1* cells, 6 kb of *cc2* DNA was replaced by 5.5 kb of *cc1* sequence, allowing the specific analysis of unique *cc2* DNA present in ectopic *cc2* insertions and minichromosomes pcc2 and pHcc2 (Catania et al., 2015). Black, red bars represent qChIP primer sites on *cc1/3* (the homologous region between *cc1* and *cc3*) and *cc2* DNA, respectively.

**Figure S2.**
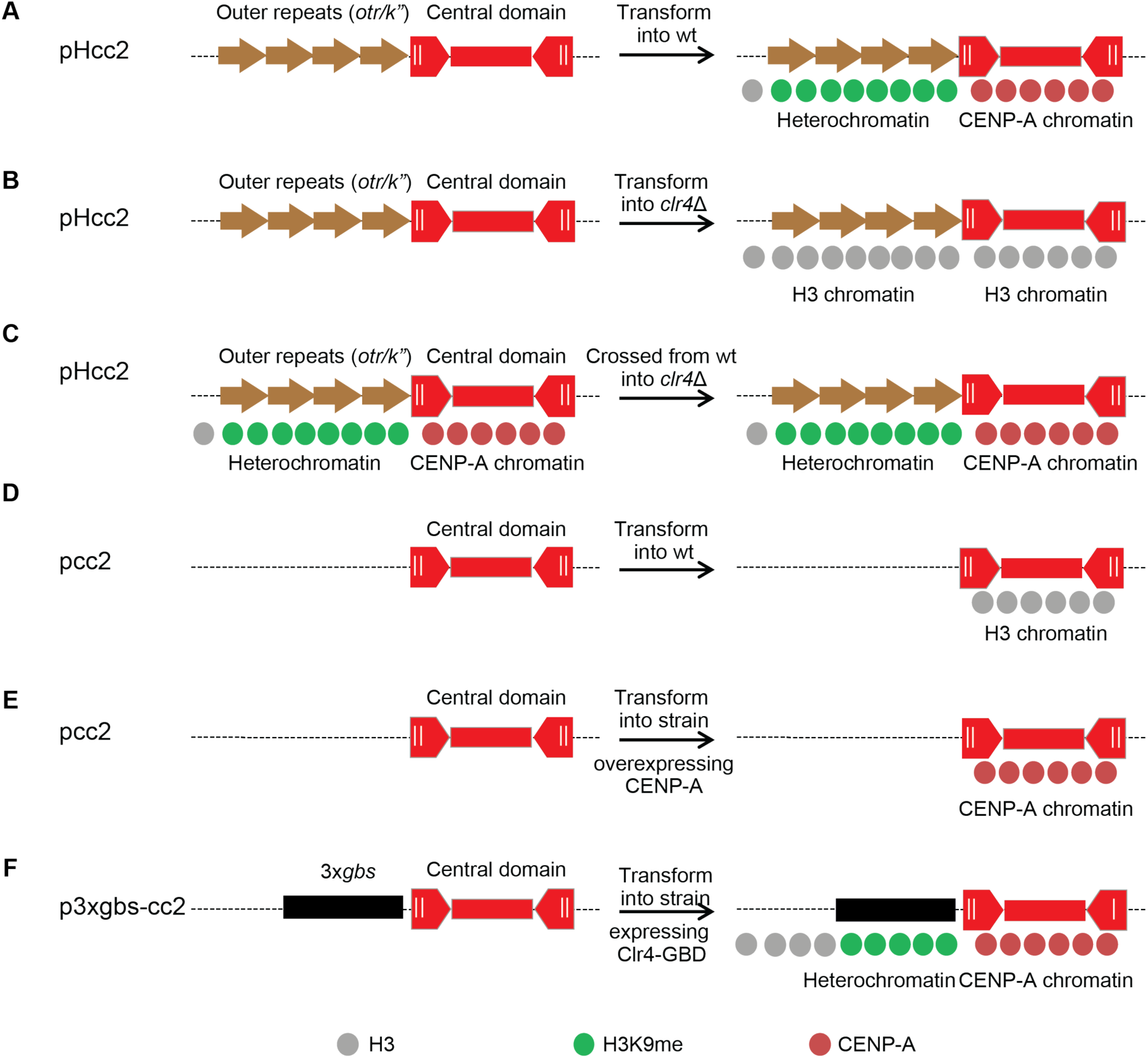
Heterochromatin is required for CENP-A^Cnp1^ chromatin establishment unless CENP-A^Cnp1^ is overexpressed. Relates to Figure 1. (A, B) pHcc2 assembles heterochromatin on outer repeats (*otr/K”*) which promotes CENP-A^Cnp1^ chromatin establishment on its central domain DNA in wt (A) cells but not in *clr4*Δ (B) cells following transformation (Folco et al., 2008). (C) Heterochromatin is not required to maintain CENP-A^Cnp1^ chromatin on central domain of pHcc2 in *clr4*Δ crossed from wt (Folco et al., 2008). (D, E) pcc2 assembles CENP-A^Cnp1^ chromatin on its central domain in CENP-A^Cnp1^ overexpressed cells (E) but not in wt CENP-A^Cnp1^ cells (D) upon transformation (Catania et al., 2015). (F) p3xgbs-cc2 assembles heterochromatin on *3xgbs* (three Gal4-binding sites) that permits CENP-A^Cnp1^ chromatin establishment on its *cc2* central domain in cells expressing Clr4-GBD (the DNA binding domain of the *S. cerevisiae* Gal4 protein) fusions following transformation. Artificial association of Clr4 with *3xgbs* bypasses the requirement for heterochromatic outer repeats (*otr/K”*) in heterochromatin assembly (Kagansky et al., 2009).

**Figure S3.**
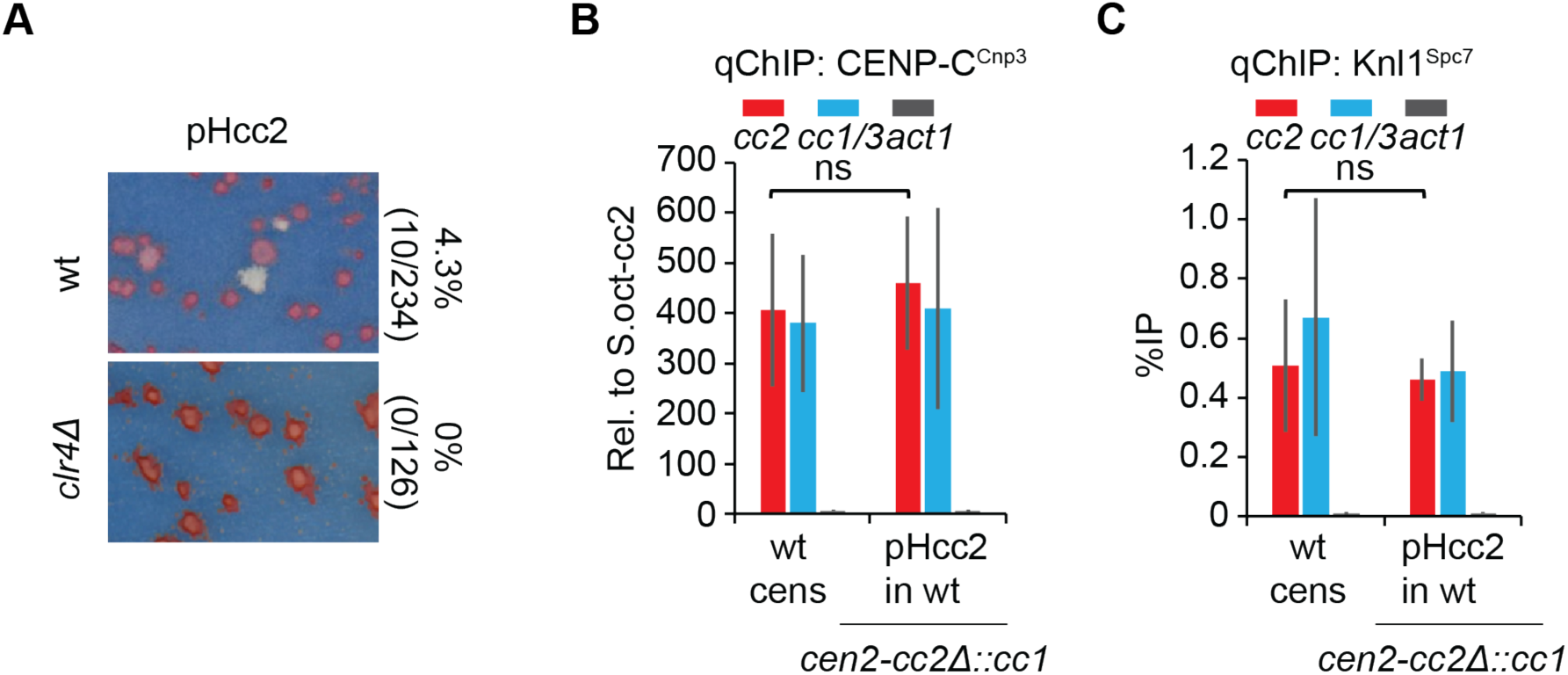
Kinetochore proteins are recruited to minichromosome pHcc2 transformed into wt cells. Relates to Figure 1. (A) Functional centromere establishment assay on pHcc2 transformed into wt and *clr4*Δ strains carrying *cc2*Δ::*cc1*. Transformants were replica plated to low adenine non-selective plate. Pale pink/white colonies indicate the presence of functional centromere since *sup3e* tRNA on pHcc2 suppresses *ade6-704* mutation within strains. Establishment frequency was calculated as the percentage of pale pink/white colonies divided by the total number of transformants (See also STAR Methods), thus 4.3%, 0% of colonies established functional centromeres on pHcc2 transformed into wt and *clr4*Δ cells, respectively. B, C) qChIP analyses for CENP-C^Cnp3^ (B), Knl1^Spc7^ (C) levels at *cc2, cc1/3* and *act1* in wt cens strain carrying endogenous *cen2-cc2* or *cen2-cc2*Δ::*cc1* strain transformed with pHcc2. %IP levels in *S. pombe* were normalized to %IP of central core from spiked-in *S. octosporus* chromatin in (B). qChIP results in (C) were reported as %IP. Data are mean ± SD (n=3). ns, no significance (Unpaired t-test).

**Figure S4.**
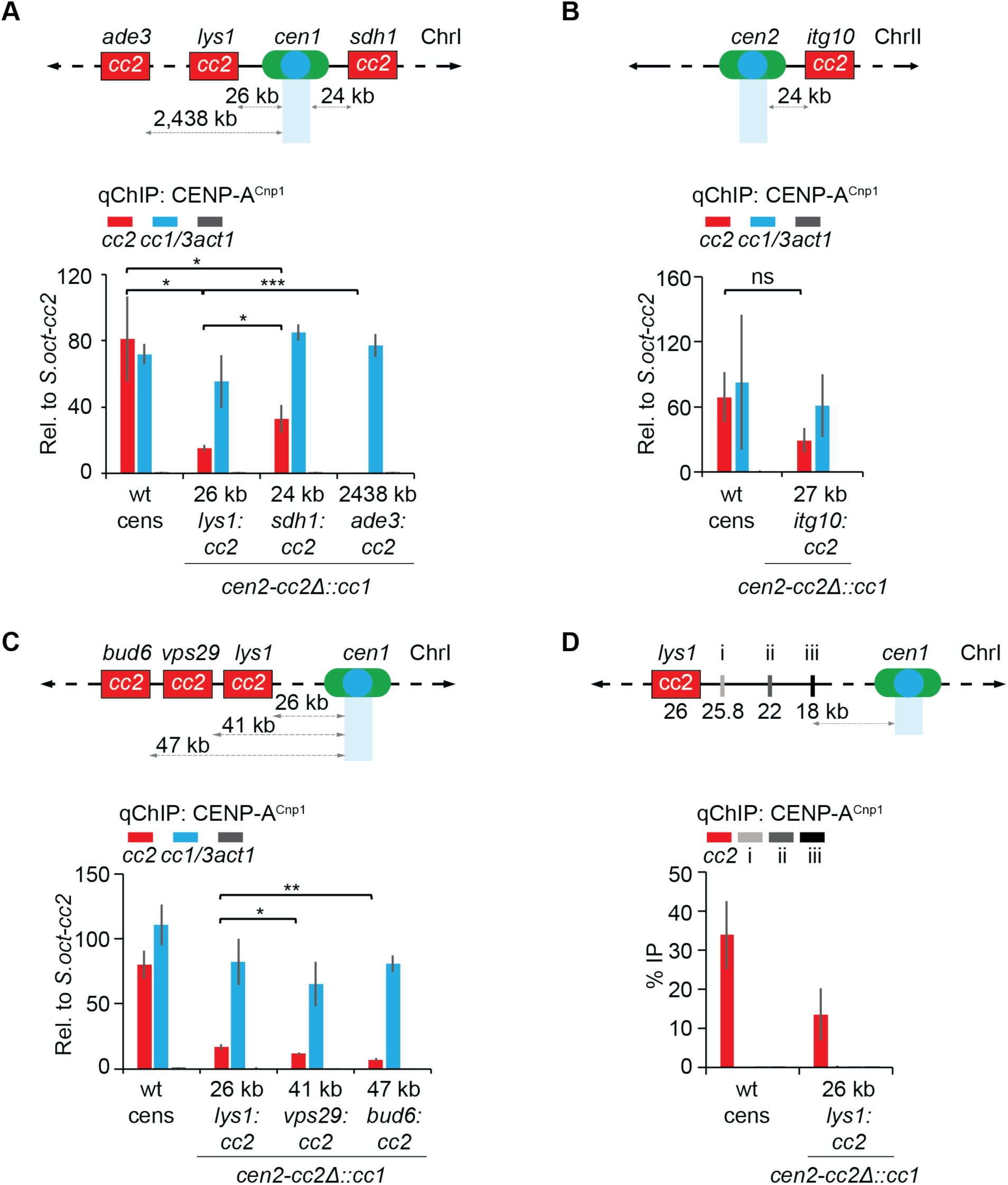
CENP-A^Cnp1^ chromatin is established on *cc2* inserted closed to centromeres. Relates to Figure 2. (A-C) qChIP analyses for CENP-A^Cnp1^ levels at *cc2, cc1/3* and *act1* in wt cens strain with *cen2-cc2* or *cc2*Δ::*cc1* strain with *lys1:cc2, sdh1:cc2*, *ade3:cc2* (A) or *itg10:cc2* (*itg10*; ChrII: 1,645,855-1,655,523; B) or *vps29:cc2* or *bud6:cc2* (C) insertion. %IP levels in *S. pombe* were normalized to %IP of central core from spiked-in *S. octosporus* chromatin. (D) qChIP analyses for CENP-A^Cnp1^ levels at three euchromatic locus between *lys1* and *cen1*: sites i, ii, iii, 25.8, 22, 18 kb from *cc1* respectively in wt cens or *cc2*Δ::*cc1* strain with *lys1:cc2.* qChIP results were reported as %IP. All qChIP data are mean ± SD (n=3). ns, no significance; *, p<0.05; **, p<0.005; ***, p<0.0005 (Unpaired t-test).

**Figure S5.**
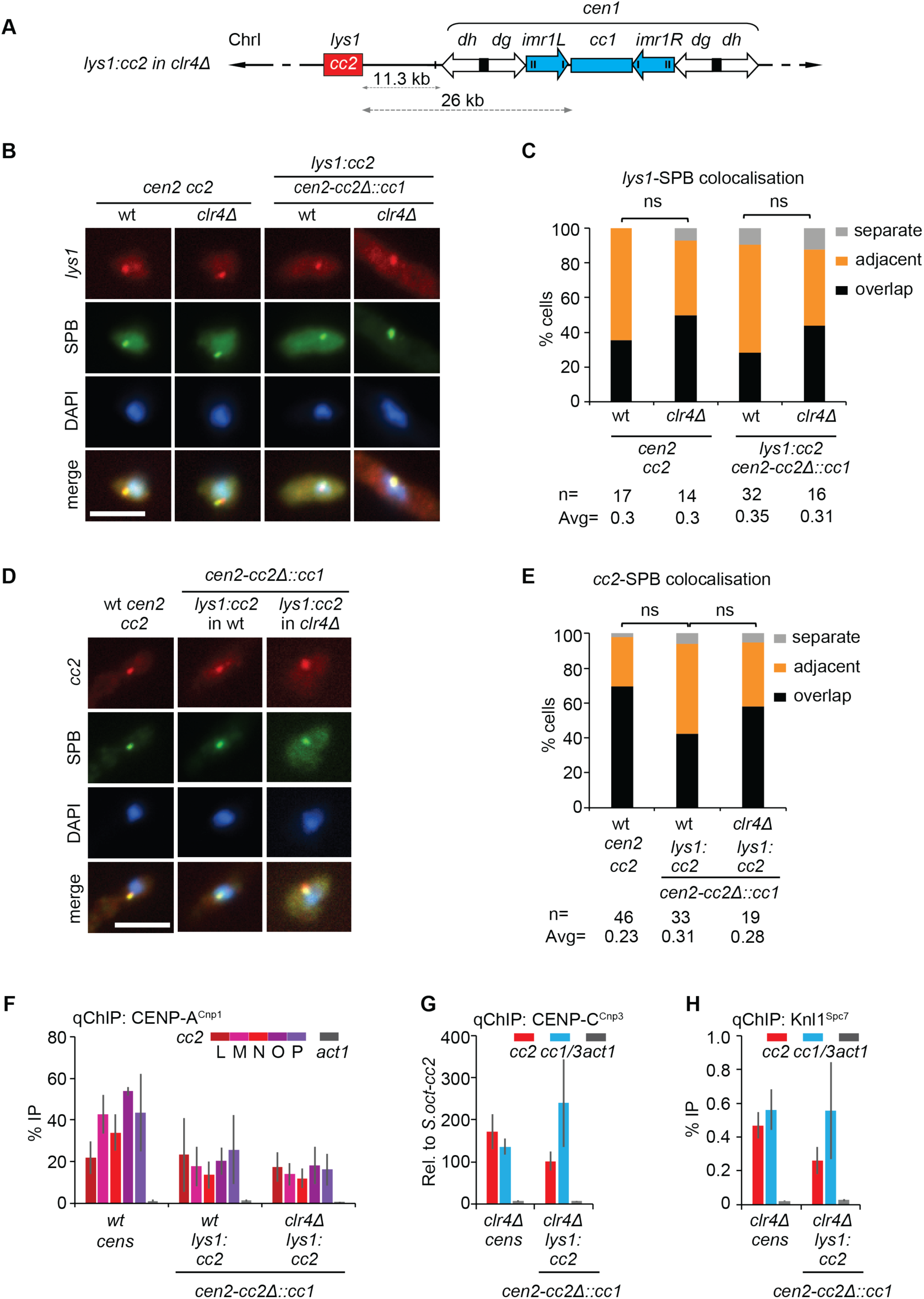
Centromeric heterochromatin is not required to establish CENP-A^Cnp1^ chromatin on *cc2* inserted close to *cen1*. Relates to Figure 2. (A) Diagram represents *lys1:cc2* insertion, 26 kb or 11.3 kb from *cc1*, *cen1 dh* repeat in heterochromatin-deficient *clr4*Δ cells following transformation, respectively. (B, D) Representative images of *lys1* (B) or *cc2* (D) DNA FISH (red), SPB location (green; anti-Cdc11) and DNA staining (blue, DAPI) in wt or *clr4*Δ strain with endogenous *cen2-cc2* or *cen2-cc2*Δ::*cc1* and *lys1:cc2*. Images were scaled as in Figure 1. Scale bar, 5 μm. (C, E) Cells were classified into three groups according to the 3D distances between *lys1* (C) or *cc2* (E) and SPB (Cdc11): overlap (≤0.3 μm), adjacent (0.3-0.5 μm) or separate (0.5-3 μm). Percentage of interphase cells (n, number analyzed from 3 independent experiments) in each category. Avg, average distance. ns, no significance (Mann-Whitney U test). F-H) qChIP analyses for CENP-A^Cnp1^ (F), CENP-C^Cnp3^ (G), Knl1^Spc7^ (H) levels at *cc2*, *cc1/3* and *act1* wt or *clr4*Δ strain with endogenous *cen2-cc2* or *cen2-cc2*Δ::*cc1* and *lys1:cc2*. %IP levels in *S. pombe* were normalized to %IP of central core from spiked-in *S. octosporus* chromatin in (G). qChIP results in (F, H) were reported as %IP. Data are mean ± SD (n=3).

**Figure S6.**
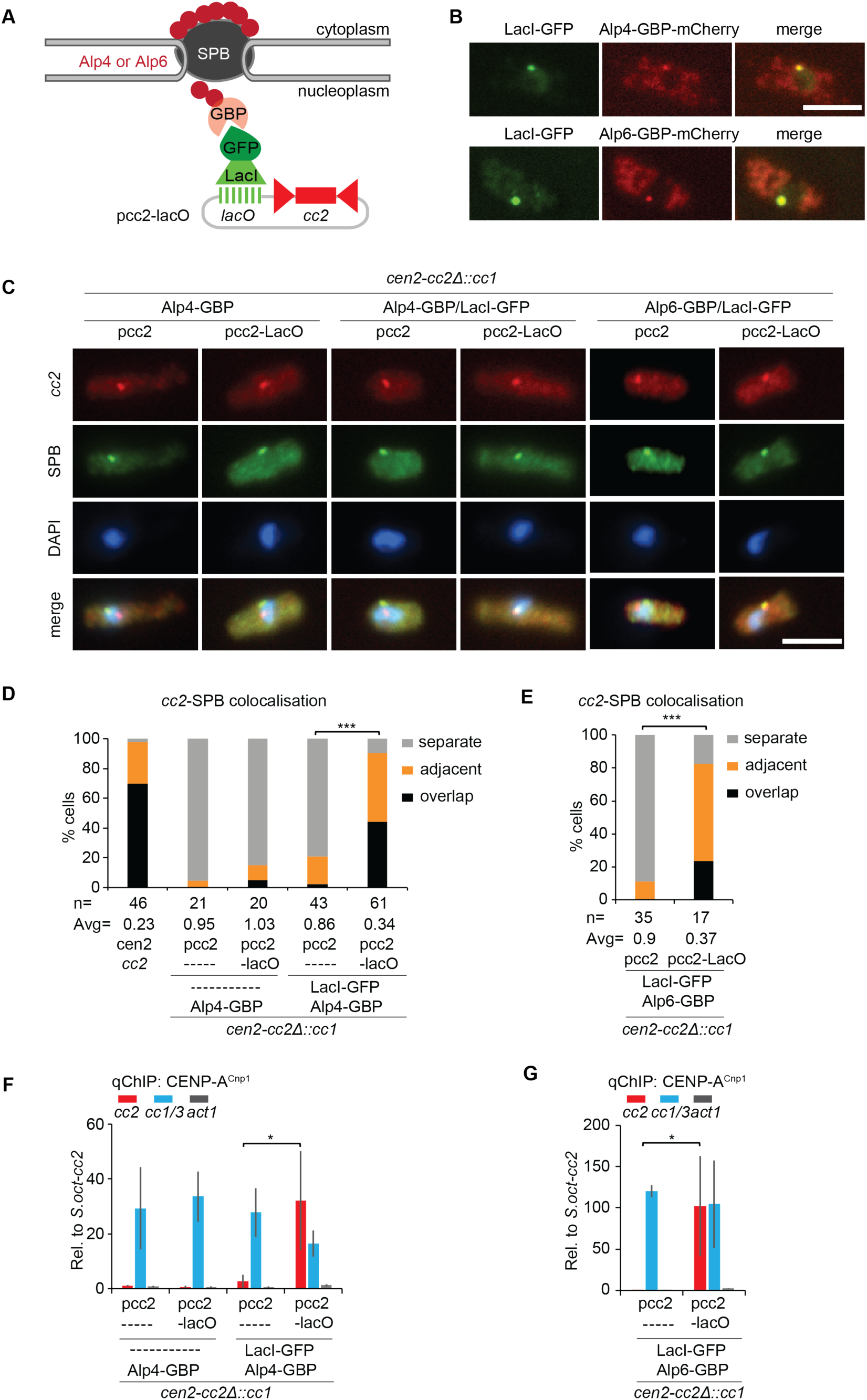
Tethering *cc2* DNA to Alp4 or Alp6 allows CENP-A^Cnp1^ incorporation, Relates to Figure 5. (A) Forced association of pcc2-lacO with Alp4 or Alp6-GBP-mCherry at SPB using same tethering system as in Figure 5. A small subset of Alp4 or Alp6 molecules (red circles) are localized to the nucleoplasmic side of SPB during interphase (Bestul et al., 2017). (B) Representative images of live cells expressing LacI-GFP and Alp4 or Alp6-GBP-mCherry. Images were scaled as in Figure 2. Scale bar, 5 μm. (C) Representative images of *cc2* DNA FISH (red), SPB location (green; anti-Cdc11) and DNA staining (blue, DAPI) in indicated strains. Images were scaled as in Figure 1. Scale bar, 5 μm. (D, E) Cells were classified into three groups according to the 3D distances between *cc2* and SPB (Cdc11): overlap (≤0.3 μm), adjacent (0.3-0.5 μm) or separate (0.5-3 μm). Percentage of interphase cells (n, number analyzed from 3 independent experiments) in each category. Avg, average distance. ***, p < 0.0001 (Mann-Whitney U test). (F, G) qChIP analyses for CENP-A^Cnp1^ levels at *cc2*, *cc1/3* and *act1* in indicated strains. %IP levels in *S. pombe* were normalized to %IP of central core from spiked-in *S. octosporus* chromatin. Data are mean ± SD (n=3-4). *, p<0.05 (Unpaired t-test).

**Figure S7.**
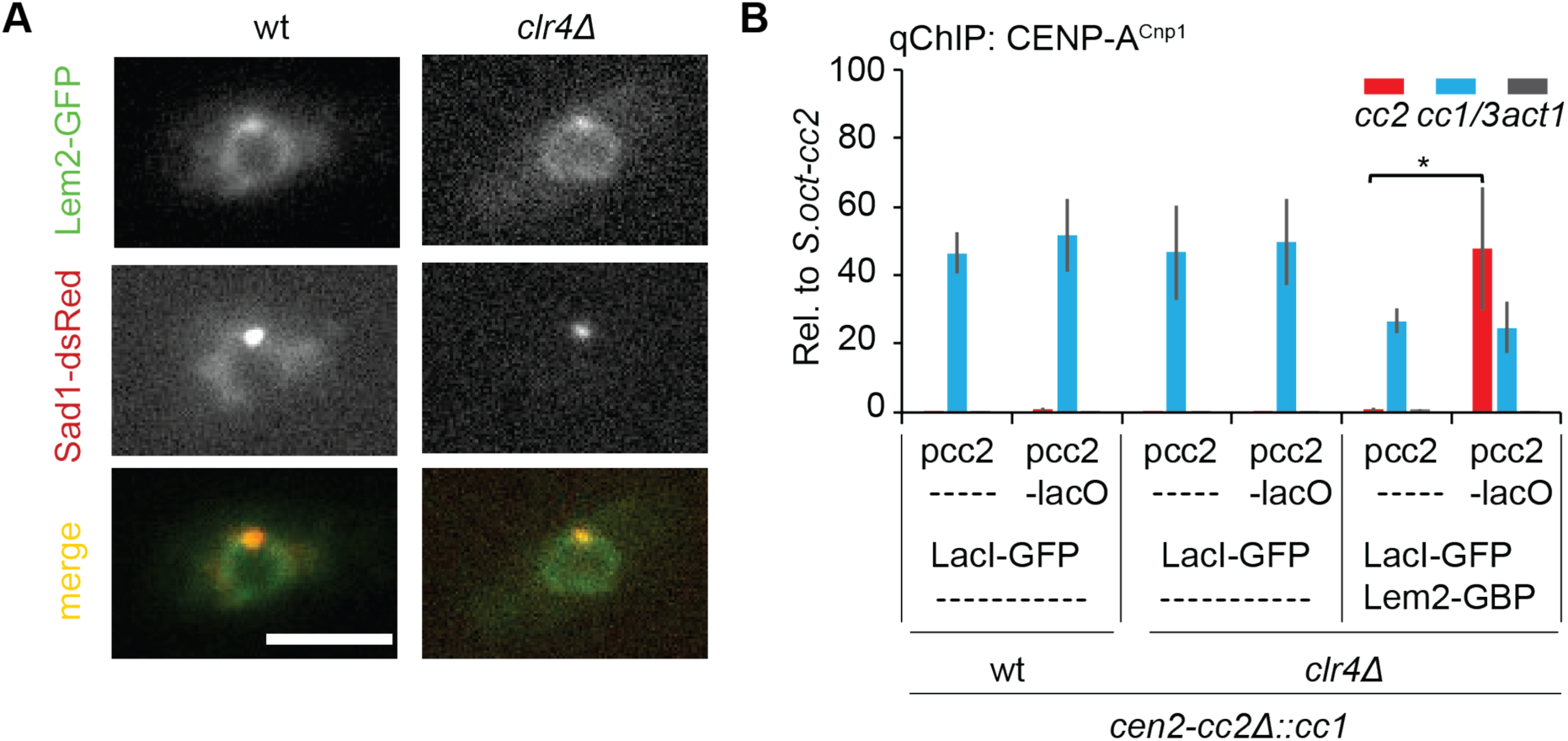
Tethering *cc2* DNA to Lem2 allows CENP-A^Cnp1^ incorporation independently of heterochromatin. Relates to Figure 5. (A) Representative images of live wt or *clr4*Δ cells expressing Lem2-GFP and Sad1-dsRed. Images were scaled as in Figure 2. Scale bar, 5 μm. (B) qChIP analyses for CENP-A^Cnp1^ levels at *cc2*, *cc1/3* and *act1* in wt or *clr4*Δ strains carrying *cen2*-*cc2*Δ::*cc1* and expressing LacI-GFP or both LacI-GFP and Lem2-GBP-mCherry transformed with pcc2 or pcc2-lacO. %IP levels in *S. pombe* were normalized to %IP of central core from spiked-in *S. octosporus* chromatin. Data are mean ± SD (n=3). *, p<0.05 (Unpaired t-test).

**Table S1.**
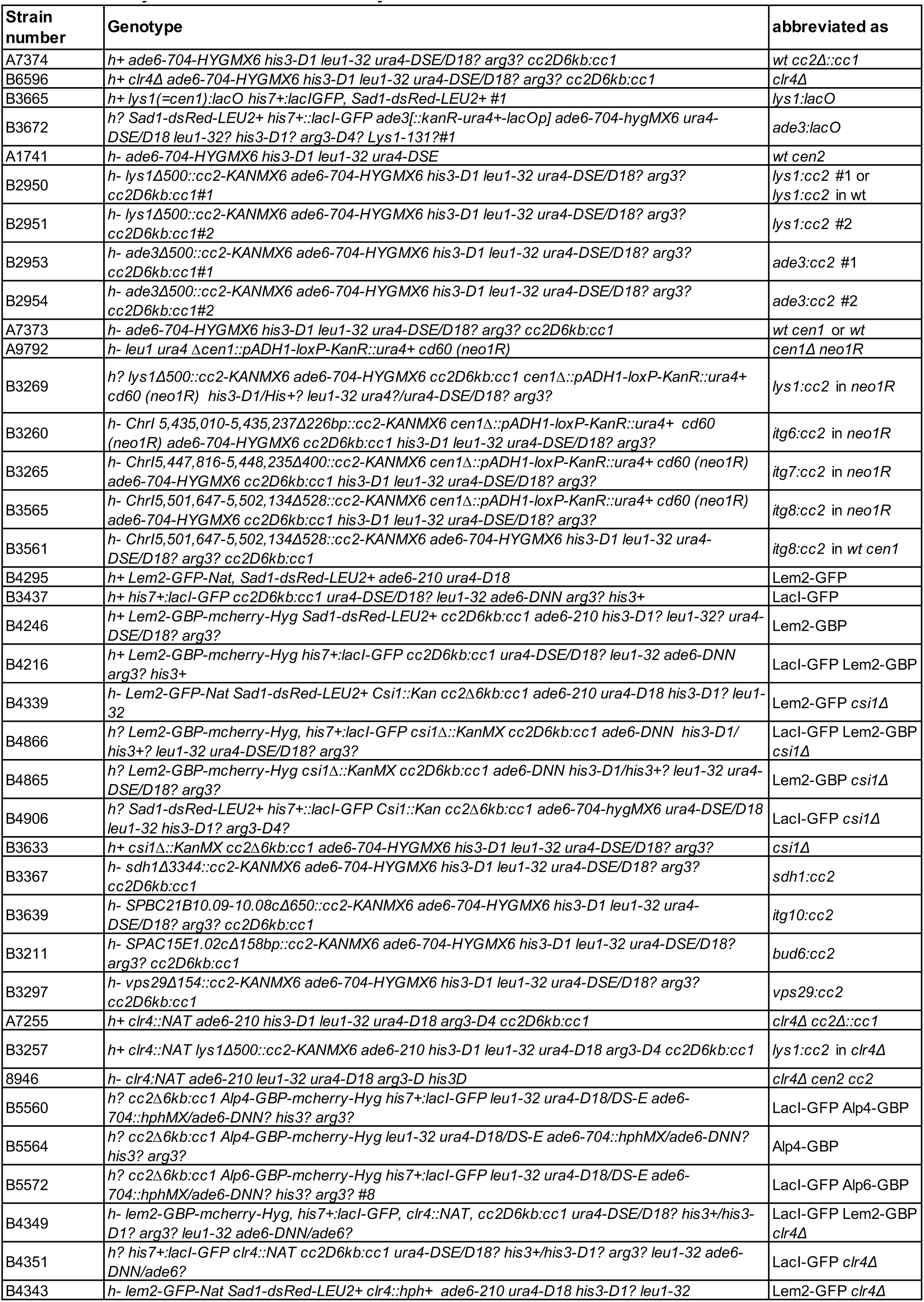
Fission yeast strains used in this study. Related to STAR Methods.

**Table S2.**
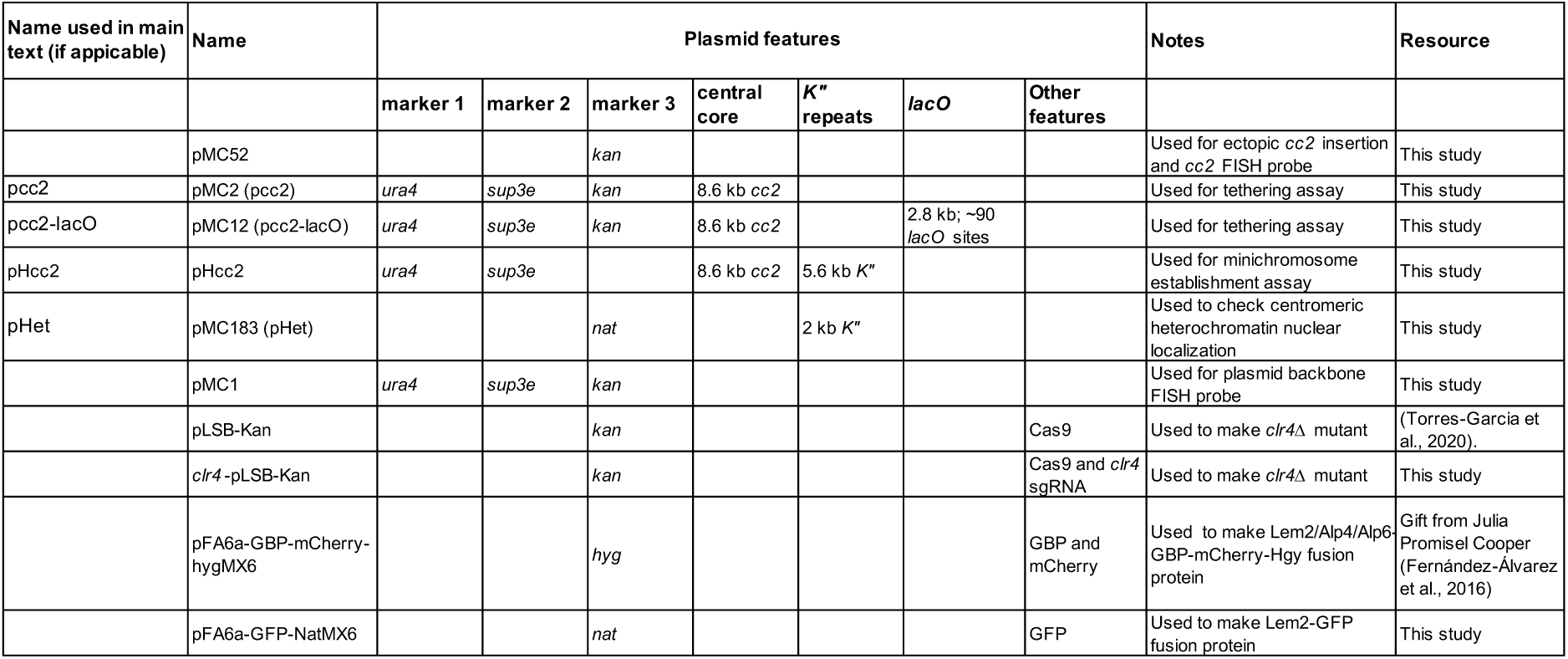
Plasmids used in this study. Related to STAR Methods.

**Table S3.**
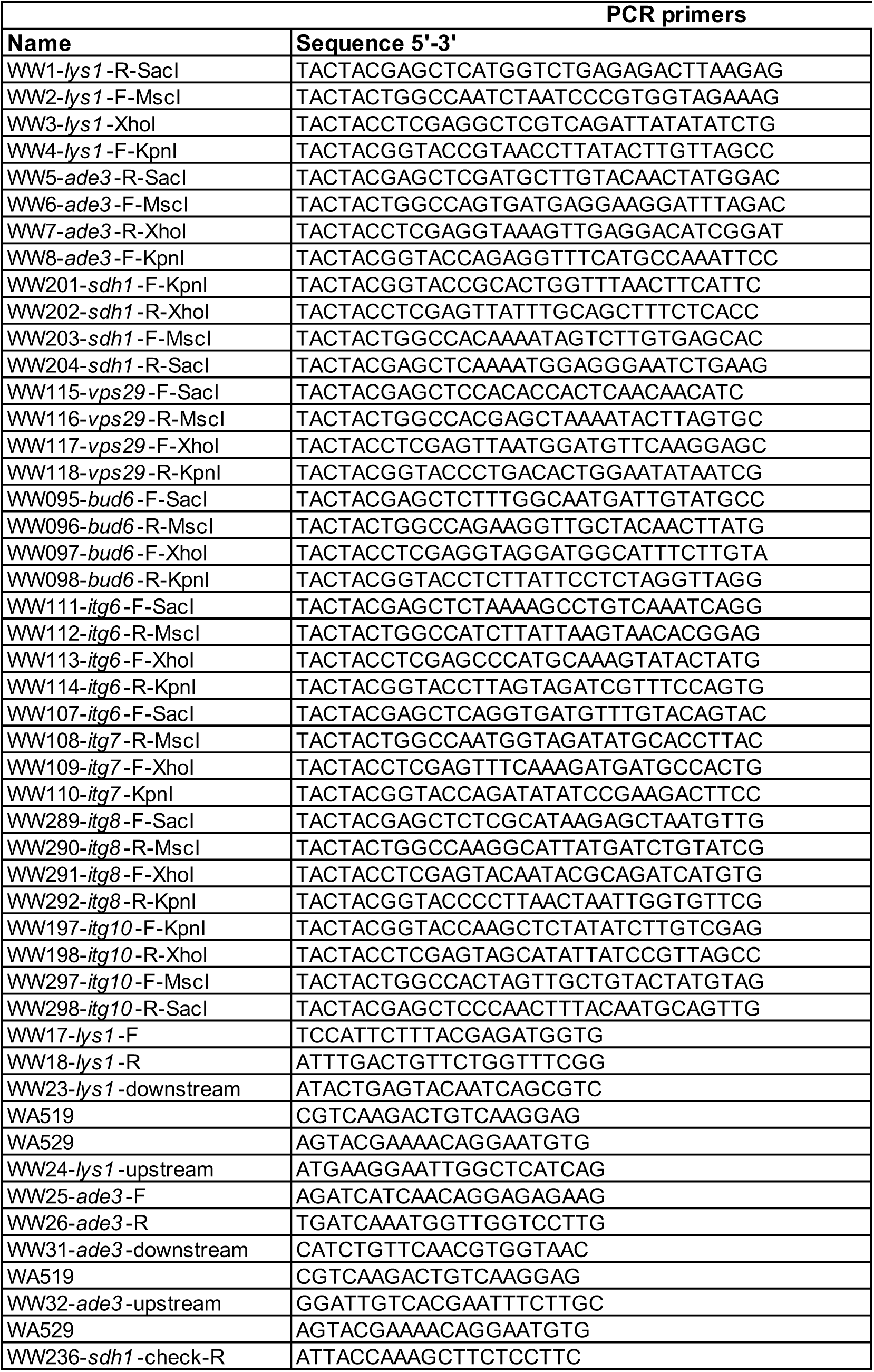
Primers used in this study. Related to STAR Methods.

